# Planting diversity begets multifaceted tree diversity in oil palm landscapes

**DOI:** 10.1101/2023.12.04.566521

**Authors:** Gustavo Brant Paterno, Fabian Brambach, Nathaly Guerrero-Ramírez, Delphine Clara Zemp, Aiza Fernanda Cantillo, Nicolò Camarretta, Carina C. M. Moura, Oliver Gailing, Johannes Ballauff, Andrea Polle, Michael Schlund, Stefan Erasmi, Najeeb Al-Amin Iddris, Watit Khokthong, Leti Sundawati, Bambang Irawan, Dirk Hölscher, Holger Kreft

## Abstract

Optimizing restoration outcomes is crucial for enhancing multifaceted diversity, resilience, and ecosystem functioning in monoculture-dominated landscapes globally. Here, we experimentally tested the performance of passive and active restoration strategies to recover taxonomic, phylogenetic, and functional diversity by establishing 52 tree islands in an oil palm landscape. Tree diversity via natural regeneration was shaped by local rather than landscape properties, with the diversity of planted tree species and tree island size driving higher multifaceted diversity. We show that large tree islands with higher initial planted diversity catalyze the recovery of multifaceted diversity at both the local and landscape level, including forest-associated species. Our results demonstrate that planted diversity begets regenerating diversity, overcoming major limitations of natural regeneration in highly modified landscapes. By elucidating the contribution of experimental, local, and landscape drivers to natural regeneration, these findings provide practical insights to make oil palm landscapes more biodiversity-friendly by enhancing functional and phylogenetic diversity within plantations.

## Main text

Globally, tropical forests are being converted into monoculture plantations, resulting in the simplification of Earth’s most complex and biodiverse ecosystems. Southeast Asian forests are known for their extraordinary biodiversity, being hotspots of plant taxonomic, phylogenetic, and functional diversity ^1,2^. However, the large-scale conversion of these forests into plantations of African oil palm (*Elaeis guineensis* Jacq.) has resulted in alarming losses of biodiversity and ecosystem functions ^3–6^. Therefore, to safeguard multifaceted tropical diversity, it is imperative to prioritize the conservation of remaining forests ^7^, improve conventional management practices ^8^, and restore human-modified landscapes aiming at biodiversity-friendly habitats that can preserve tropical biodiversity outside of protected areas^9,10^.

Ecological restoration can contribute to bending the curve of biodiversity loss by enhancing biodiversity in degraded ecosystems while improving ecosystem functioning, resilience, and human well-being ^11–13^. Recent research highlights the importance of viewing restoration approaches as a continuum, ranging from minimal intervention, i.e. passive (e.g., natural regeneration and management practices to promote natural regeneration), to intensive assistance required for ecosystem recovery, i.e. active (e.g., tree planting and major soil modification) ^14^. Recognizing this continuum is crucial because passive and active restoration methods can synergistically complement each other. Natural regeneration is a restoration strategy that can be applied at large scales ^15,16^ and has been shown to be effective in restoring degraded ecosystems in temperate and tropical regions ^12,17,18^, including agroecosystems ^19^. Active restoration, for instance, via planting tree islands is a promising restoration strategy promoting rapid forest regeneration in tropical regions ^20–22^.

Recent findings suggest that tree islands established through active tree planting can enhance multi-taxa species diversity and ecosystem multifunctionality within large-scale oil palm plantations ^10,23^. However, increasing also functional and phylogenetic diversity of restored ecosystems is an important step towards improving ecosystem functioning, resilience to environmental disturbances or disease outbreaks, adaptation to changing conditions such as climate change or invasive species, and nature’s contribution to people ^24–27^. Despite this, studies assessing restoration of multiple facets of biodiversity are still rare ^28–30^.

Enhancing multifaceted diversity via tree planting in combination with natural regeneration likely relies on local and landscape context ^31^. Natural regeneration can be difficult to predict and may be influenced by local and landscape drivers (**Extended Data Figure 1, Table S1**) ^15,32^. Active tree planting can promote natural regeneration by improving local conditions and facilitating the recruitment of trees and shrubs ^33,34^, potentially accelerating the overall recovery of phylogenetic and functional diversity ^30^. Planting a higher diversity of trees can thus enhance vegetation structural complexity, i.e. the three-dimensional distribution of plants in the ecosystem, affecting biotic interactions within and across trophic levels ^35^ and promoting plant growth and ecosystem resilience ^36,37^. Tree island size influences on regenerating diversity mainly arise from island biogeography theory, i.e. larger islands increase species diversity through higher colonization and lower extinction rates ^38^. Larger tree islands also promote plant colonization by reducing environmental stress and edge effects while simultaneously attracting more seed dispersers ^21,39^. Furthermore, in human-modified landscapes, adverse soil characteristics can limit seedling establishment ^40^. The landscape context may also influence the local species diversity within tree islands ^41^. For example, areas closer to forests receive more seeds and experience higher plant colonization rates ^42^ and trees scattered in the agricultural matrix act as stepping stones for seed dispersers and as pollen sources ^43^. Understanding the relative importance of local vs. landscape factors contributing to natural regeneration is therefore crucial to predicting successional trajectories ^32^ and guiding more effective restoration efforts on degraded lands worldwide ^15,31^.

Here, we assessed the local and landscape factors affecting the recovery of taxonomic, phylogenetic, and functional tree/woody species diversity within 52 tree islands established in a large-scale oil palm plantation located in Sumatra, Indonesia. The quadratic island plots vary in planted tree diversity (0, 1, 2, 3, 6 species - with 0 corresponding to natural regeneration only) and size (25 m² to 1,600 m²) (**Fig S8**). Weeding and grazing were stopped to allow for natural regeneration within the islands (EFForTS-BEE [Ecological and socio-economic functions of tropical lowland rainforest transformation systems: biodiversity enrichment experiment] ^44^. We tested if local multifaceted diversity of the regenerating trees is driven by local conditions (i.e., experimental treatments and soil properties), landscape context (i.e., distance to the nearest forest and density of scattered trees), or both using structural equation modeling (see **Extended Data Figure 1** and **Table S1** for more details on the mechanistic framework and predictions). We hypothesized that the influence of the experimental treatments on tree regeneration acts directly and indirectly via vegetation structure (i.e. vegetation complexity and tree dominance) (**Extended Data Figure Fig 1**). Specifically, we expected that larger tree islands with higher richness of planted trees would directly increase recruiting diversity by providing a more heterogeneous environment and thereby wider niche space, and reduced competition due to complementarity in resource uses between different species. Additionally, we expected vegetation structural complexity provided by planted trees to indirectly influence recruiting diversity by providing contrasting habitats that allow the establishment of tree species varying in their functional strategies. We also anticipated increased tree dominance to promote natural regeneration by reducing understory cover of light-dependent species and competition with oil palms. Lastly, we predicted that tree islands with higher soil quality and located closer to propagule sources (forest patches and scattered trees) would exhibit greater recruiting diversity due to improved establishment conditions and reduced dispersal limitations, respectively (**Extended Data Figure 1; Table S1**).

**Figure 1.**
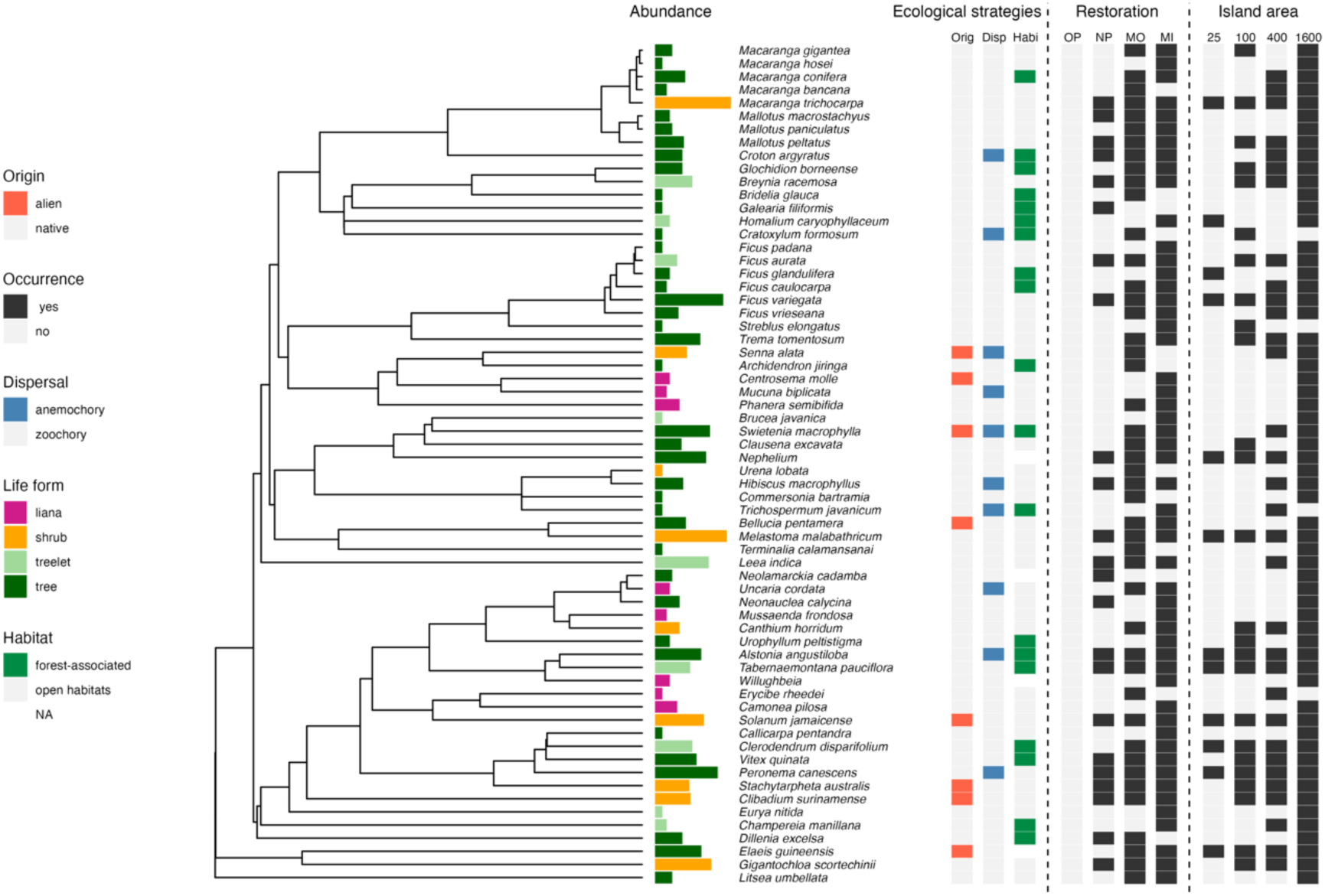
Phylogenetic relationships, abundances, ecological strategies, and occupancy of regenerating species. Bar colors represent life forms (liana, shrub, treelet, and tree) of all 64 woody species regenerating in the EFForTS-BEE experiment in Sumatra, Indonesia. Bar length represents species’ total abundance across plots (log-scale), ranging between 1-1218 individuals. One morphospecies (climber DMF0061) was not included in the phylogenetic tree. Species origin (alien vs native), dispersal mode (anemochory vs zoochory), and habitat (forest vs open habitat) are highlighted in red, blue, and green, respectively. Black rectangles indicate occurrence in different restoration treatments: OP: oil palm; NP: no planting; MO: monoculture (only one planted tree species); MI: mixture plantations (2-6 planted tree species); and tree islands with different areas: 25 m^2^, 100 m^2^, 400 m^2^, and 1600 m^2^.

We find that, already six years after establishment, tree islands hosted a high multifaceted diversity of recruiting trees, which differed greatly from the impoverished oil-palm landscapes under conventional management. Specifically, we documented a diverse range of woody species established through natural regeneration, representing 4386 woody plant individuals belonging to 65 species from 32 plant families and reflecting different plant ecological strategies (**Fig 1**, see **Table S2** and **Fig S1-2** for an overview of functional trait combinations). The five most represented families, in terms of number of species, were Euphorbiaceae (N = 9 species), Moraceae (N = 7), Rubiaceae (N = 6), Fabaceae (N = 5), and Lamiaceae (N = 4). Regenerating woody species represented various plant ecological strategies, including trees (N = 39), treelets (N = 9), shrubs (N = 9), lianas (N = 9), bamboo (N=1) (**Fig 1**, **Fig S1**). Most species (N = 41) were associated with secondary forests and disturbed habitats, while one-third were forest-associated species (N = 19). Forest-associated species had higher maximum height and lower specific leaf area (SLA) than species associated with open habitats (**Fig S2**). Zoochory was the most prevalent dispersal syndrome, accounting for approximately 85% of all individuals (N = 3732) and 85% of species (N = 56). The primary dispersers of recruiting species were birds, bats, and mammals. While high levels of zoochory are expected in tropical forests, our results elucidate the need to protect functioning seed disperser communities to ensure successful restoration.

Tree islands host a range of rare, common, and abundant species, with rarer species mainly located on larger islands. Overall, abundance was highly correlated with occupancy (R^2^: 0.84, **Fig S1d**), with shrubs and trees among the locally most abundant and widespread species. However, some species were locally abundant but rare across plots (e.g. *Swietenia macrophylla*, n = 172 individuals / 4 plots), while others were locally rare but widespread across plots (e.g. *Solanum jamaicense*, n = 97 ind. / 29 plots). Further, fifteen species were only registered once (singletons). A small fraction of species was non-native to Sumatra (N = 8), representing less than 10% of regenerating plant individuals (**Fig S1**).

### Planting diversity begets multifaceted gamma diversity in larger tree islands

At the landscape level, woody species multifaceted diversity showed stronger increases in islands with planted trees (either single species or mixtures) (**Fig 2 a-c**) and larger tree islands (**Fig 2 d-f**), compared with no species regenerating in conventionally managed plots (mean = 0). Differences between single and mixture plantings were more pronounced for taxonomic and phylogenetic diversity and for Hill diversity of order q=0 (**Fig 2a-c**, **Fig S3**). Larger tree islands (≥ 400 m^2^) with mixed planting hosted most singletons, with 40% of species solely registered in the largest tree islands (i.e. 1600m^2^, **Fig 2g**). Larger islands also supported a much higher abundance of trees (**Fig 2g**, **Fig S11**), ensuring not only the presence of rare and forest-associated species but also common and abundant species (see **Fig S3** for details). At the tree island scale, species richness ranged from 0 to 24, while the average number of species per plot was 7.2 ± 6.4 SD (**Fig S11**). Island area and restoration treatment jointly explained a large amount of the variance in observed alpha diversity (q=1) (R^2^_tax_ = 0.70, R^2^_phy_ = 0.72, and R^2^_fun_= 0.70) (**Table S3**), with a consistent positive effect of the island area across diversity facets and between observed and standardized diversity (i.e. accounting for sampling effort through coverage-based standardization), and even stronger for observed Hill diversity (q=0) (**Fig S5-6**). Tree islands with mixed species showed, on average, the highest observed alpha Hill diversity (q=1) (**Extended Data Figure 2**). Yet, besides a higher average compared to tree islands with only one planted tree species, when considered as a categorical variable, the effect of tree islands with mixed species was only significant for functional diversity of recruited trees (**Extended Data Figure 2**).

**Figure 2.**
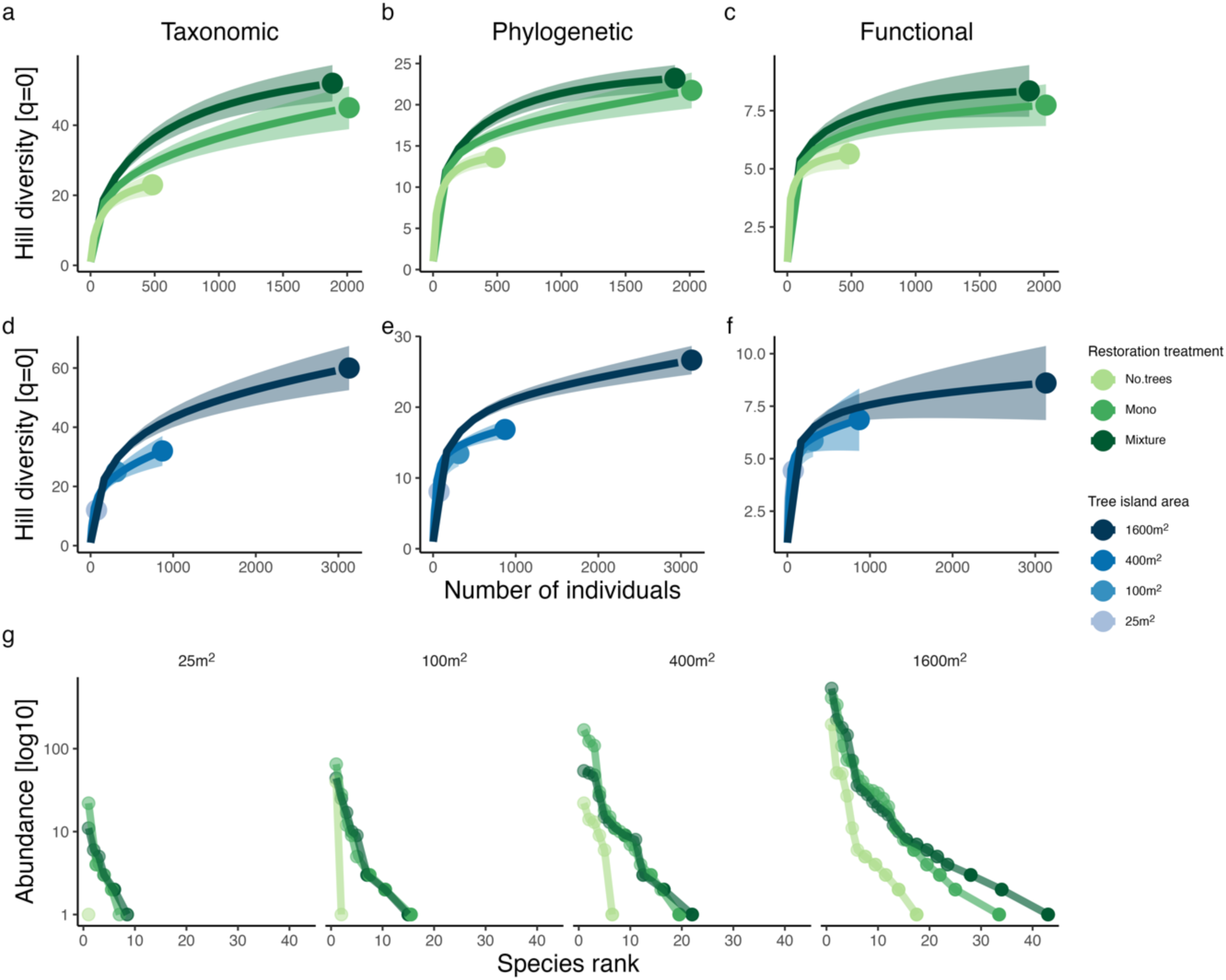
Mixed planting and larger tree islands enhance multifaceted diversity at the landscape scale (gamma diversity). **a-f,** Sample-size-based rarefaction curves for taxonomic, phylogenetic, and functional Hill diversity [q=0] across restoration treatments **(a, b, c**) and across tree island areas (**d, e, f**). Solid lines represent rarefaction curves based on a complete census of the tree islands. Solid points represent observed Hill diversity. Error bars represent 95% confidence intervals from bootstrapping (N = 500 randomizations). **g**, Rank-abundance curves for different restoration treatments (no planting, monoculture, and mixture) and tree island area sizes (25 m^2^, 100 m^2^, 400 m^2^, and 1600 m^2^). Species abundances were log10 transformed.

Increased multifaceted diversity of recruiting trees as a result of tree planting, particularly at landscape level, supports growing evidence demonstrating that tree planting can accelerate natural regeneration compared to no planting (i.e. natural regeneration only) ^30,33,34,45,46^. Planted trees, through facilitation, can promote natural regeneration by alleviating environmental stress and improving local conditions for recruiting species that otherwise cannot establish and survive in disturbed areas ^47,48^. Facilitation can further promote species regeneration through plant-animal interactions (e.g. secondary seed dispersal) ^49^ and tends to be stronger and more frequent for rare than for common species ^50^. In addition, how established tree species modify environmental conditions and resource availability (i.e. impact niche) can promote recruiting species with contrasting growth and survival requirements (i.e. requirement niche) ^51,52^, leading to an increased functional diversity of recruiting trees. Recruited individuals with contrasting ecological strategies may also benefit from reduced competition due to niche complementarity in mixed plantings ^53^, improved tree growth and performance ^54^, and greater stability in growth ^55^ and survival.

At local scales, based on the SEM model, diversity of planted tree species had a direct positive effect on the observed taxonomic (Std. beta = 0.21, p = 0.033; 95% CI [0.02, 0.41]), functional (Std. beta = 0.270, p = 0.016; 95% CI [0.05, 0.49]), and phylogenetic diversity (Std. beta = 0.176, p = 0.085; 95% CI [−0.03, 0.38], **Fig 3abc**), although the positive effect was not statistically significant for phylogenetic diversity (p = 0.084). Our results provide needed evidence of the impact of overstory tree diversity on long-term ecosystem dynamics via regenerating trees ^56,57^. Existing studies are mostly observational (e.g., based on large forest inventory data) ^58,59^, with experimental studies focusing mostly on the influence of tree diversity on biodiversity of associated taxa (herbaceous vegetation or other taxa) and multiple ecosystem functions ^60–62^. In one of the few studies that experimentally tested the effects of planted tree species diversity on woody recruitment diversity, planted tree diversity did not influence recruiting species richness after five years in an experiment in Costa Rica ^57^. Differences between studies may be likely related to the closer proximity to existing forests and minimal soil degradation at their study sites. Therefore, our results provide unique experimental evidence of the pivotal role of how planting diversity begets regenerating diversity in cash-crop dominated landscapes, where the islands can mitigate dispersal and establishment limitations of the surrounding landscape. Further, the results of our experiment have clear implications for restoration ecology and suggest that initially planted diversity, through a positive feedback loop on regenerating diversity, can maintain and promote multifaceted diversity over time, potentially enhancing ecosystem functioning and resilience of restored areas to future conditions.

### Effects on diversity beyond passive sampling

Island area emerged as the overall most important driver of recruiting diversity, with tree island area affecting positive multifaceted woody species diversity, directly and indirectly, through enhanced tree dominance (**Fig 3abc**, see **Table S4** and **Fig S7** in the supplementary material). The strong and consistent positive effect of tree island area on multifaceted diversity is consistent with theoretical and empirical evidence suggesting that species diversity may increase with sampling area due to passive sampling ^38,63^. However, tree island area also had a strong positive effect when controlling for sampling effort (e.g., coverage-based standardized diversity) (**Fig S4**), supporting additional ecological mechanisms playing a role beyond passive sampling. For example, for standardized Hill diversity (q=1), the positive effect of tree island area shifted from direct to indirect via enhanced tree dominance, leading to increased taxonomic, phylogenetic, and functional alpha diversity (**Fig 3def**). The positive effect of tree dominance on recruiting diversity can be explained by higher tree cover providing better habitat quality via effects on light and microclimate ^22,39^, being more attractive (than open areas) for seed dispersers such as birds, bats, and mammals ^20,21,39^. Larger tree islands have a more heterogeneous environment and thereby provide wider niche space. Reduced competition with oil palm in large tree islands may also have played a role, with previous results showing that planted trees growing near oil palms experience reduced growth ^64^ and increased mortality. Increased tree dominance was also correlated with reduced cover of invasive alien plants (e.g., *Clidemia hirta*), increased litter cover, higher water infiltration ^65^ and more stable microclimatic conditions ^66^, which may increase seedling survival and diversity. Our results highlight that increases in restoration area may have disproportionately larger positive effects on biodiversity gains and sustainable populations than would be expected from passive sampling alone.

### Local drivers play a major role on large-scale industrial plantation and landscape drivers

Soil properties (PC1, reflecting higher soil C, N, and lower soil compaction) had a significant and positive effect on taxonomic (Std. beta = 0.30, p = 0.003; 95% CI [0.11, 0.49]), phylogenetic (Std. beta = 0.28, p = 0.007; 95% CI [0.08, 0.47]) and functional diversity of woody species (Std. beta = 0.30, p = 0.006; 95% CI [0.09, 0.52]). In contrast, soil properties reflecting phosphorus (P) and pH (PC2: Soil properties) had a positive but non-significant effect on the taxonomic (p = 0.2960), phylogenetic (p = 0.2392), or functional diversity (p = 0.1823) (**Fig 3abc; S7**). Oil palm plantations can reduce soil fertility and increase soil compaction over time ^67^, which may limit the establishment of small seedlings attempting to regenerate ^40^. Thus, in conventional oil palm plantations, improving soil fertility and reducing soil compaction may be an effective way to reduce establishment limitations of recruiting woody species, with potential cascading effects on biodiversity. Particularly, when considering that extensive areas under oil palm cultivation are approaching replanting age (∼25 years), leading to further soil degradation and constraining the possibilities for future multifaceted restoration efforts.

Contrary to our initial expectations, when considering local and the landscape contexts simultaneously, the distance to the nearest forest, and the abundance of scattered trees surrounding tree islands had little or no detectable effect on recruitment diversity **(Fig 3**, **Fig S7**). Previous studies have often reported that proximity to primary forests or increased forest cover increases biodiversity by reducing dispersal limitation ^42,68^. However, other studies have found little or no landscape effect on restoration outcomes ^69,70^. The conditions under which landscape context is more or less relevant are still debated ^15,31^, may vary with taxa and scale and early versus later successional stages ^41^ or when multiple drivers are considered simultaneously. The lack of landscape influence in our study can be explained by the absence of primary forests and low secondary forest cover in our study region (∼4%) (**Extended Data Figure 1**). The overall limited amount of surrounding forest patches, typically observed in tropical large-scale agricultural landscapes, may hinder the influence of forest distance on recruitment diversity. Additionally, abundant generalist animal species that thrive in oil palm landscapes and facilitate long-distance seed dispersal may also explain the lack of effect of surrounding forests. Disturbance-tolerant animal species can become primary seed dispersers when larger vertebrates are absent from the landscape ^71^. For example, long-tailed macaques (*Macaca fascicularis*) and common palm civets (*Paradoxurus hermaphroditus*) are known to facilitate seed dispersal of several plant species ^72^ and both species occur at our site. Therefore, a priority in cash-crop landscapes should be to conserve a high coverage of forests and other landscape components, such as scattered trees, which can facilitate positive restoration outcomes. Moreover, future work is needed to understand the role of disturbance-tolerant species in promoting natural regeneration in agricultural tropical landscapes.

**Figure 3.**
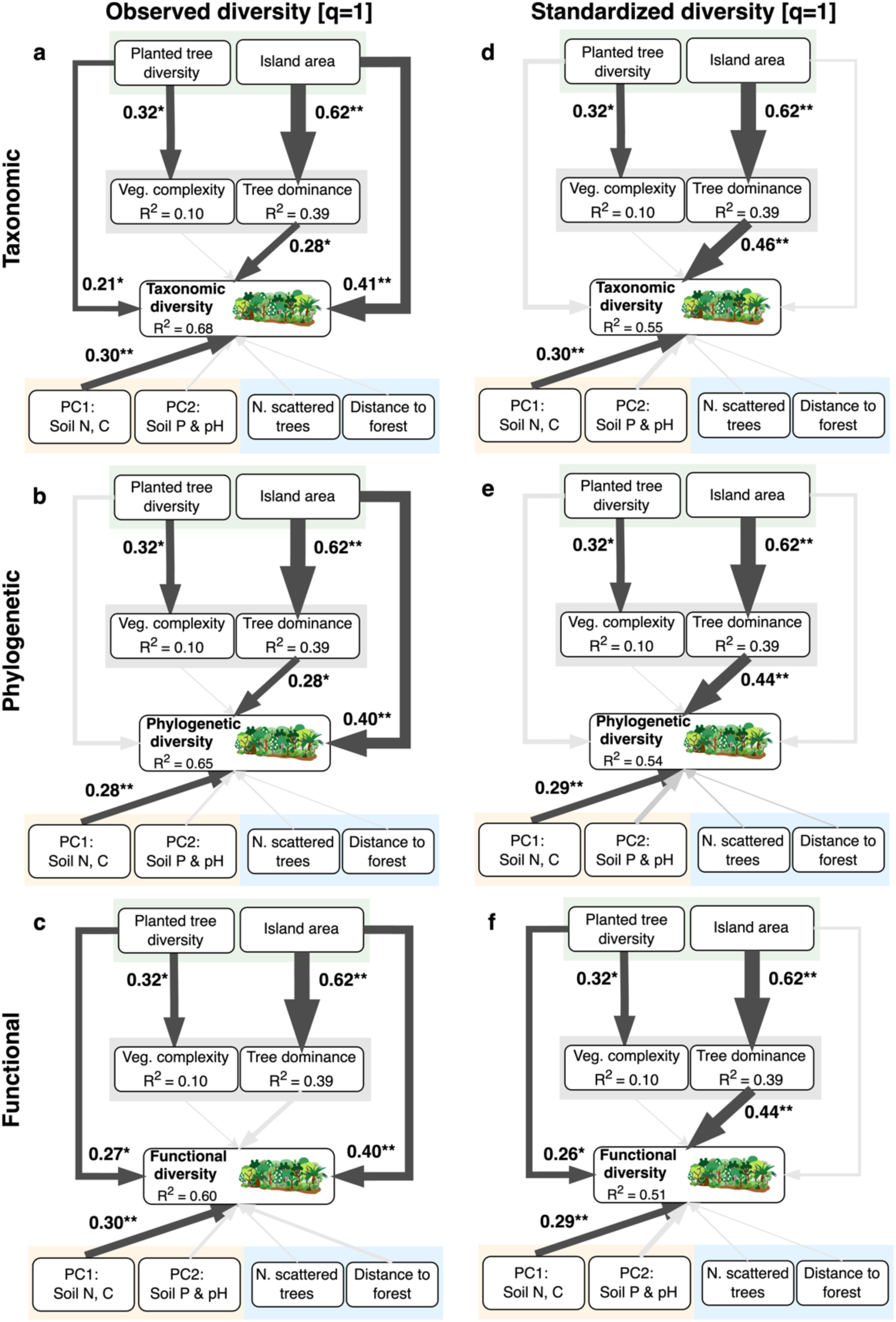
Local and landscape drivers of multifaceted diversity of recruiting woody species. Results from Piecewise Structural Equation Models explaining the main drivers of observed (**a, b, c**) and standardized (e, f, g) Hill diversity [q=1] of the regenerating plant community in EFForTS-BEE, Sumatra, Indonesia. Black arrows indicate positive effects. Gray arrows are non-significant paths (P value < 0.05). The thickness of paths reflects standardized model coefficients (in bold). * p<0.05 ** p<0.01 *** p<0.001. Diversity estimates were standardized to the minimum sample coverage (SC = 0.75) across tree islands extrapolated to double reference size.

### Implications for restoration ecology

Our results suggest that strategic choices in restoration planning can successfully guide restoration outcomes, confirming the viability of the tree island approach to catalyze the recovery of plant diversity in agricultural lands ^21,39^. Moreover, our results suggest that different restoration strategies can be used to maximize different facets of diversity at different scales. For example, larger tree islands appear crucial to maximize the taxonomic and phylogenetic diversity at the landscape scale by supporting rare and forest-associated species (**Fig 2; Fig S3**). On the other hand, to maximize functional diversity within tree islands, it may be required to increase the number of planted tree species to promote species recruitment with contrasting ecological strategies (**Fig 3c**; **Extended Data Figure 2; Fig S2**). These results highlight how incorporating functional and phylogenetic dimensions into restoration ecology provides a more holistic perspective, improving our understanding of restoration outcomes.

### Conclusion

Recognizing the urgent need to advance our scientific knowledge on ecosystem restoration is imperative for addressing the threats posed by the large-scale conversion of tropical forests into monoculture plantations worldwide and for achieving the goals of the UN Decade of Ecosystem Restoration^73^. Through a highly controlled, long-term field experiment, we demonstrate that initial restoration decisions, such as the diversity of planted species and restoration area, can have long-lasting effects on multifaceted diversity in the natural regeneration at plot and landscape scales. Our results provide strong evidence that combining diverse tree planting with natural regeneration in large tree islands is an effective strategy to catalyze the recovery of taxonomic, phylogenetic, and functional diversity in conventional palm plantations.

## Materials and Methods

### Study site

This study was conducted inside a 140 ha oil palm plantation (PT. Humusindo Makmur Sejati, 01.95° S and 103.25° E) in Jambi province, Sumatra. The study region is dominated by oil palm plantations (83%). In contrast, other land use types, such as fallow (6%), secondary forest (4%), and rubber plantations (2%), are present in patches distributed across the landscape ^74^. The region has a tropical climate (Af, according to the Köppen classification) with a mean annual precipitation of 2577 mm and a mean annual temperature of 26.1 °C. The rainy season is between October and March, while the months with the lowest precipitation (i.e. ∼ 100 mm/month) are between June and September.

### Experimental design

The EFForTS-BEE experiment was established in December 2013 (see ref. ^10,44^ for further details) and is part of a global network of biodiversity experiments (TreeDivNet) ^75^ and of a Collaborative Research Center between Indonesia and Germany (CRC990 EFForTS, http://www.uni-goettingen.de/crc990) ^76^. EFForTS-BEE consists of four partitions with varying plot sizes (5 m x 5 m, 10 m x 10 m, 20 m x 20 m, and 40 m x 40 m) and diversity of planted tree species (0, 1, 2, 3, and 6). The experiment comprises 56 plots, including four control plots with oil palm management as usual and 52 experimental tree islands (**Fig S8**). Tree islands with varying tree species diversities and compositions were established randomly within the oil palm plantation with a minimum distance of 85 m between them. The experimental treatments, i.e. diversity and identity of the planted species and island size, varied following a random partition design ^77^.

Six native multi-purpose trees were selected for planting, including three tree species mainly grown for fruits (*Parkia speciosa* Hassk., Fabaceae; *Archidendron jiringa* (Jack) I.C.Nielsen, Fabaceae; *Durio zibethinus* L., Malvaceae), two species used for timber (*Peronema canescens* Jack, Lamiaceae; *Rubroshorea leprosula* (Miq.) P.S.Ashton & J.Heck. [previously known by its synonym: *Shorea leprosula* Miq.], Dipterocarpaceae), and one species which produces natural latex (*Dyera polyphylla* (Miq.) Steenis, Apocynaceae) ^64^. Before tree planting, selected oil palms were removed (by approximately 40%) to enhance the environmental conditions for the planted trees in the experimental plots, except in control plots and the 5 m x 5 m plots ^78^. Trees were planted in a 2 m x 2 m grid in alternating rows in a north-south direction in December 2013. On mixed-species plots, trees of the same species were planted as far away as possible from one another. The number of trees planted varied depending on the plot size, with six trees on the 5 m x 5 m plots, 25 trees on the 10 m × 10 m plots, 100 trees on the 20 m × 20 m plots, and 400 trees on the 40 m × 40 m plots. In total, 6354 trees were planted in the experiment. The management of the oil palm plantation surrounding the experimental plots includes the application of fertilizers, manual weeding of the understory and occasional application of herbicides. To enhance tree survival, a single dose of plant inorganic fertilization was applied at the time of tree planting. From then on, applying fertilizer, pesticides, or herbicides stopped entirely in the tree islands (excluding control plots). Manual weeding around the planted trees was carried out only for the first two years. See ref. ^44^ for further details on the experimental design.

### Data collection

#### Plant inventory

In the 56 experimental plots, all free-standing woody plants, including trees, shrubs, lianas, and bamboos, with a length of ≥ 130 cm, were recorded in 2020. The minimum length was tested with a 1.3-m-long stick, which was also used to find the point to measure the diameter. For each morphospecies, one voucher specimen with two duplicates was collected after taking field photographs. Afterwards, the field photographs and specimens were used to identify the species of all collected plants with the help of relevant floras (Flora Malesiana, Tree Flora of Malaya, Flora of the Malay Peninsula) and databases (POWO, Digital Flora of Indonesia), as well as taxonomic monographs and revisions, where necessary. *Clidemia hirta*, a common invasive shrub occurring in disturbed sites, was not included in the inventory because it was too numerous. Instead, the cover of *Clidemia hirta* within a 5 m x 5 m subplot was estimated for each experimental plot ^10^. Cover of *Clidemia hirta* was on average 32% ± 18.6 SD. Plant taxonomic classification was harmonized according to the World Checklist of Vascular Plants (WCVP) ^79^.

#### Phylogenetic tree

We built a phylogenetic tree that included all 68 species in this study (planted tree species plus regenerating species). As backbone, we used the mega-tree provided by ^80^, containing the global angiosperm phylogenetic tree developed by ref. ^81^ and the pteridophyte tree from ^82^. We built our study tree with the R-package “V.PhyloMaker2” ^83^. Twenty-six species were missing from the global phylogeny. These were included in the study tree using scenario 3, which binds new tips at halfway of the family branch, when the genus is missing, or at the basal node of the genus, when the new tip belongs to an existing genus at the backbone phylogeny ^80^. This scenario is the most recommended for community analysis ^84^.

#### Plant functional traits

Key vegetative and dispersal traits were collated from open databases. A literature search was performed to acquire dispersal syndrome and plant life form, while maximum plant height (m) and life forms were obtained from Flora Malesiana (https://floramalesiana.org/) and specialized botanical literature. Specific leaf area (mm^2^/mg) and wood density (gm/cm^3^) were obtained from several sources, including the TRY database ^85^, Global Inventory of Floras and Traits (GIFT) ^86^ and dataset provided by refs. ^87,88^. For a subset of eleven species, we used wood density information at the genus level from ref. ^87^ global database. Species origin (alien vs. native to Sumatra) was obtained from the World Checklist of Vascular Plants (https://powo.science.kew.org/) ^79^, complemented with regional botanical literature. Plant dispersal mode was obtained from regional and specialized botanical literature. See the supplementary Materials for details on data sources and plant trait data. We collated a database on species life forms, dispersal syndrome, and maximum height for all species. Species-level data were missing for wood density (26%) and specific leaf area (50%). To deal with missing data, we used Rphylopars, a maximum likelihood method that utilizes a phylogeny and a sparse trait matrix that reconstructs ancestral states and imputes missing values ^89^. This approach has been demonstrated to be the most accurate for handling missing data for continuous traits ^90,91^. We used the R-package “Rphylopars” ^89^ to impute missing values for specific leaf area (SLA) and wood density (WD). Diversity estimates using raw and imputed datasets were highly correlated and did not affect our primary analyses or trait distributions. Therefore, only the results using imputed data are presented.

#### Vegetation structure and complexity

We acquired 12 different variables representing various aspects of vegetation structure ^10^, including four indicators derived from terrestrial laser scanning ^92,93^, measured in September to October of 2016: (1) stand structural complexity index (SSCI), (2) the mean fractal dimension index (MeanFRAC), (3) the effective number of layers (ENL), and (4) the understory complexity index (UCI). Using hemispherical photo and drone-based photogrammetry taken in September to October 2016 ^74^, we calculated (5) canopy gap fraction and canopy cover, that was partitioned as (6) oil palm cover and (7) tree cover; and (8) oil palm density. We conducted ground-based assessment of (9) tree density, (10) understorey vegetation cover, (11) litter cover and (12) litter depth. We then standardized all the variables to zero mean and unit variance and applied a Principal Component Analysis (PCA). The first main component (PC1) represented a gradient from open to dense and complex vegetation (“structural complexity”) and the second component (-PC2) represented a gradient from oil palm to tree dominance (“tree dominance), which were subsequently used in the statistical analysis (**Fig S9**). More information on the structural variables and the PCA can be found in the supplementary material (**Fig S9**) and in ref.^10^.

#### Soil fertility

In December 2016, an initial soil sampling was conducted during the rainy season. Three soil cores were extracted from each of the 56 research plots, resulting in 168 samples. Soil samples were collected from management zones in control plots to avoid additional variability. The samples were sieved, cleaned of roots and litter, and then freeze-dried. Total carbon (C), nitrogen (N), and plant available phosphorus (P) were analyzed using Bray and Kurtz method^94^. Soil pH was measured in a KCl suspension according to ISO 10390 standard at the University of Göttingen in Germany. More information on the soil variables can be found in ref. ^95^.

#### Landscape metrics

Two landscape-level metrics were computed: the distance of each experimental plot to the nearest forest patch (Dist. Forest) and the number of isolated trees in a 100 m buffer from the experimental plots (scattered trees). For the nearest forest patch, forest areas were classified from ortho-aerial photographs at 0.1-m resolution using fixed-wing drone (Aero M, 3D Robotics, USA) equipped with red, green, blue and near infrared cameras (Canon PowerShot SX260 HS, Japan) ^74^. The orthophotographs were then processed using supervised classification and post-processing steps to generate a 1-m resolution land use map ^74^. Secondary forest patches were converted to polygons and the distance from each plot coordinate to the nearest forest polygon was measured using the near function in ArcGIS (version 10.4).

Scattered trees were manually annotated using very high-resolution (5 cm) airborne true color orthophotos and a Light Detection and Ranging (LiDAR) derived canopy height model (CHM) at 1 m resolution. Data was collected in January 2020 over the studied area. The number of scattered trees in a 100 m buffer was calculated in a GIS environment (QGIS v.3.22.8) using zonal statistics. We also computed the number of scattered trees using different buffers: 100, 200, and 500 m from the plot’s center. A buffer of 100 m was used in the main analyses because this was the scale with the strongest effect on recruiting diversity.

### Diversity Estimation and Statistical Analysis

#### Estimating diversity with Hill numbers

Hill-Chao numbers were used to estimate the taxonomic, phylogenetic, and functional alpha diversity of the regenerating woody species for each plot ^96^. Hill-Chao numbers are a family of diversity measures that capture the effective number of species present in a community ^97,98^. This approach provides a unified framework for estimating diversity across multiple scales and biodiversity facets ^99,100^. To account for sampling bias, besides observed diversity, we also estimated coverage-based diversity with rarefaction/extrapolation of Hill numbers ^101^. We used the maximum sampling coverage following the approach suggested by ^102^. This methodology ensures that diversity estimates are robust and unbiased by accounting for potential sampling bias in the data and standardizing diversities to the same unit: the effective number of equally abundant species (taxonomic diversity), the effective number of equally divergent lineages (phylogenetic diversity), and the effective number of equally distinct functional species (functional diversity) ^96^. For functional diversity, we computed the dissimilarities in traits (SLA: specific leaf area, WD: woody density, MH: maximum height, and LF: life form) using a modification of Gower distance ^103^, which computes a multi-trait dissimilarity matrix with a quasi-identical contribution of individual traits ^104^. For computing the dissimilarity matrix, we used the “gawdis” function in the R-package “gawdis” ^104^. For phylogenetic diversity, we used the phylogenetic distance between species to compute a pairwise phylogenetic distance matrix.

We estimated Hill diversities using the three q-orders 0, 1, and 2. When q=0, the Hill number represents the species richness of the community. When q=1, the Hill number represents the exponential of the Shannon index (Shannon-Hill) and considers species abundances without favoring neither rare nor common species. When q=2, the Hill number represents the inverse of the Simpson index (Simpson-Hill) and reflects the diversity of the most abundant species. To calculate observed and coverage-based standardized taxonomic, phylogenetic, and functional diversity, we used the R package iNEXT.3D ^96^.

### Statistical modeling

Linear models were used to test whether enriching oil palm plantations with native trees promotes the recovery of plant taxonomic, phylogenetic, and functional diversity through natural regeneration (Analysis 1). Hill diversity (taxonomic, phylogenetic, functional) was used as response variables against two additive predictors: restoration treatment (oil palm, no planting, tree planting with single species, and mixed tree planting) and island area (25 m2, 100 m2, 400 m2, 1600 m2) (See Supplementary Material for details on model specification). Hill diversity and island area were log-transformed to improve model residuals. We tested multiple pairwise comparisons across treatment levels through least-squares means with P-values adjusted by the Tukey method. Pairwise comparisons were performed with the R-package “emmeans” ^105^.

To understand the main internal and external drivers of natural regeneration in enriched oil palm plantations (Analysis 2), we used confirmatory path analysis ^106,107^. Piecewise structural equation models (SEM) were fitted using the R-package “piecewiseSEM” ^108^. We built a conceptual model (**Extended Data Figure 1, Table S1**) describing the hypothesized causal paths between the diversity of regenerating woody species and four groups of predictors: (i) the experimental treatments (planted tree diversity, island area), (ii) vegetation structure (PC1_veg_: vegetation complexity and -PC2_veg_: tree dominance), (iii) soil properties (PC1_soil_: soil carbon, nitrogen, and bulk density; PC2_soil_: soil phosphorus, pH), and (iv) landscape context (distance to the nearest forest patch, number of scattered trees in a 100 m radius) (**Fig 1**). See the supplementary material for details on the Principal Component Analysis for vegetation complexity and soil fertility and the distribution of each variable included in the model (**Table S5; Fig S9**).

The goodness of fit was tested with Fisher’s C statistics ^107^, and potentially missing paths were evaluated with the d-separation test ^108^. Indirect effects for planted tree diversity and island area were calculated by multiplying standardized coefficients along pathways in the model. Confidence intervals for direct, indirect, and total effects for each predictor were calculated in the R-package “semEff” ^109^ using nonparametric bootstrap with 1000 randomizations.

#### Model validation

Model assumptions, including normality of residuals, homogeneity of variance, and multicollinearity, were checked by visual inspection of residuals and statistical tests using the R-package “performance” ^110^. Additionally, we tested model residuals for spatial autocorrelation using Moran’s I autocorrelation coefficient in the “ape” R-package ^111^. To check for spatial autocorrelation of the response variables (taxonomic, phylogenetic, and functional diversity), we used spatial correlograms based on Moran’s I autocorrelation coefficient, which indicated no spatial autocorrelation for any of the response variables (**Fig S10**).

## Acknowledgements

We thank PT Humusindo for granting us access to and use of their properties. We thank Eduard Januarlin Siahaan, Krisman Hakim Dalimunthe, Edo Mauliarta, Dian Muh. Fauzan, and M. Ihsan for their support in field campaigns. We thank Martin Ehbrecht and Dominik Seidel for contributing data on vegetation structural complexity. This study was funded by the Deutsche Forschungsgemeinschaft (DFG, German Research Foundation) – project number 192626868 – SFB 990 in the framework of the collaborative German–Indonesian research project CRC990. This research was conducted under the research permits: 100/SIP/IV/FR/2/2023 (Gustavo B. Paterno), 46/E5/E5.4lS|P.EXT/2019 (Fabian Brambach), 58/SIP/IV/FR/10/2021 and 20/SIP/IV/FR/1/2022 (Carina C. M. Moura), 1092/FRP/SM/VIII/2015 and 1487/FRP/SM/KI VI/2016 (Watit Khokthong). The EFForTS-BEE experiment is part of TreeDivNet, a global network of tree diversity experiments (https://treedivnet.ugent.be/).

## Author Contributions Statement

G.B.P., F.B., N.G.R., D.C.Z., H.K., and D.H. conceived this study. G.B.P., F.B., D.C.Z., N.C., C.C.M.M., J.B., A.P., W.K., and M.S. contributed data. G.B.P. developed the R code and performed formal analysis. G.B.P. wrote the original draft with contributions from N.G.R, F.B., D.C.Z., H.K., and D.H. G.B.P., L.S., B.I., D.H., and H.K. were responsible for the project administration. H.K., D.H., L.S., B.I., O.G., A.P., S.E. were responsible for funding acquisition. All authors contributed to editing and reviewing the final version of the manuscript.

**Correspondence and requests** for materials should be addressed to Gustavo Brant Paterno

## Data availability statement

The data that support the findings of this study will be made available in GRO.data repository.

## Code availability statement

The code supporting all findings of this study will be made available in GRO.data repository.

---

**Extended Data Figure 1.**
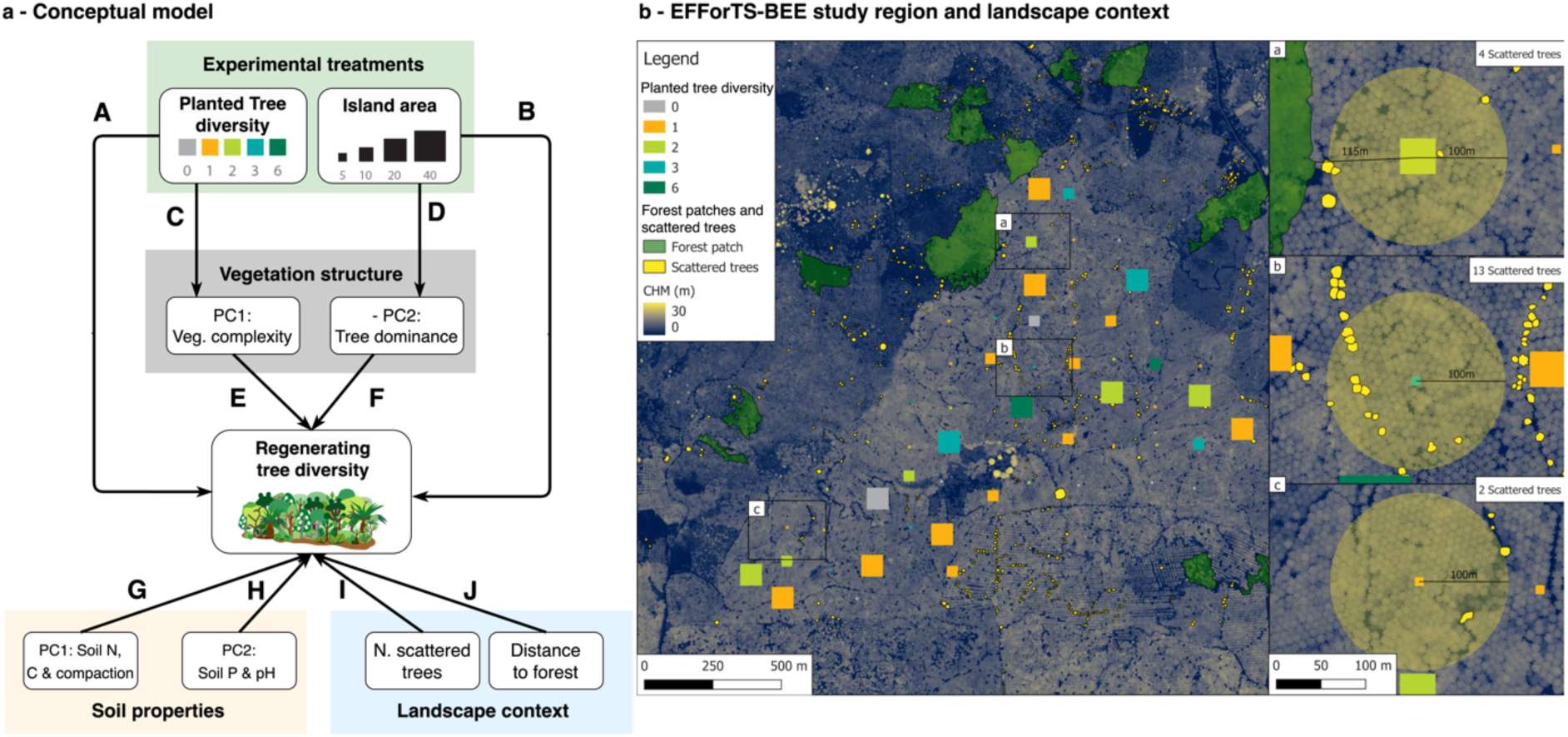
Conceptual model describing the main drivers influencing the diversity of natural regeneration. **a,** Colored rectangles represent groups of drivers: experimental treatments, vegetation structure, soil properties, and landscape context (distance to nearest forest and number of scattered trees). White round boxes represent measured variables. Arrows represent hypothesized unidirectional causal links between variables. Potentially missing links were evaluated as part of the model fit through d-separation test. Given that a previous analysis showed no effect between experimental treatments and soil properties ^95^, these paths were not included. See supplementary material for details. Mechanistic relationships for each path code (bold letters in upper case) are described in Table S1. **b**, Landscape map illustrating the spatial distribution of tree islands, secondary forest patches (larger than 0.5 ha)^74^, and scattered trees. Buffer area (100m) used to calculate the number of scattered trees is illustrated in subpanels a-c. Canopy height model (CHM) was derived from airborne LiDAR scans.

**Extended Data Figure 2.**
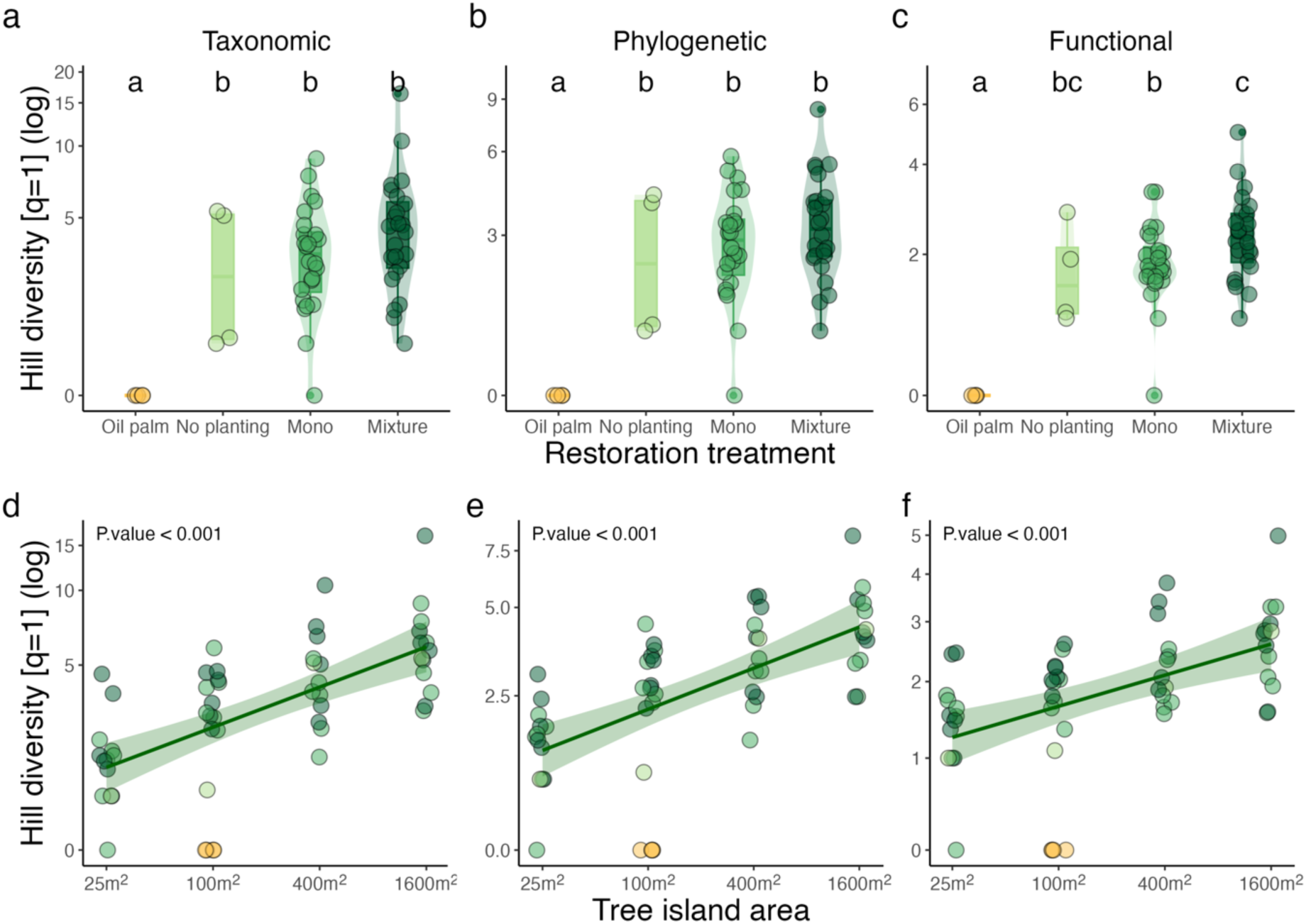
Experimental effects on multifaceted alpha diversity of recruiting woody species. **a-f,** Relationship between observed taxonomic, phylogenetic, and functional Hill diversity (q=1) (log-scale) against restoration treatment (a, b, c) and tree island area (d, e, f). Yellow points represent conventional oil palm monoculture, while green points represent tree islands. Darker green represents higher planted tree diversity, and light green represents plots with no planted trees. In boxes, horizontal lines indicate the median, and the upper and lower hinges indicate 75 % and 25 % quantiles, respectively. Letters represent pairwise comparisons across treatment levels through Least-Squares Means with p-values adjusted by the Tukey method.

