## Supplementary material for "Planting diversity begets multifaceted tree diversity in oil palm landscapes"

### This File contains:

Supplementary Text

5 Tables

11 Figures

**Supplementary Text**

**Statistical protocol for analysis 1: “Does enriching oil-palm plantations with native trees promote the recovery of taxonomic, phylogenetic, and functional diversity through natural regeneration?”**

*Modeling approach*

We fitted a linear model (estimated using ordinary least squares) to predict the diversity of regenerating species with island area and restoration. Standardized parameters were obtained by fitting the model on a standardized dataset version. 95% Confidence Intervals (CIs) and p-values were computed using a Wald t-distribution approximation.

*Response variables*

- tax_hill: Taxonomic hill diversity
- phy_hill: Phylogenetic hill diversity
- fun_hill: Functional hill diversity

*Explanatory variables*

- island_area: Island area ($m^2$) | Range: 25 $m^2$ - 1600 $m^2$
- restoration: Conventional oil palm, no planting, mono (single species planting), mixed (2-6 planted species)

*Model formula:*

```
mod <- stats::lm(hill ~ island_area + restoration, data = datset)
```

*Model validation*

For each linear model, we:

- Used a QQ-plot of residuals to check for normality
- Tested model residuals for spatial autocorrelation using Moran’s I coefficient
- Tested for homogeneity of variance
- Inspected residuals for multicollinearity
- Checked for influential observations (outliers) based on a composite outlier score (Lüdecke et al., 2021)

**Statistical protocol for analysis 2: “What are the main internal and external drivers of natural regeneration in enriched oil-palm agroforestry systems?”**

*Data pre-processing before analysis*

- Dataset was cropped to tree islands only by excluding the four control plots (conventional oil palm)
- All numeric predictors were centered (substracting the mean) and scaled (dividing centered values by the standard deviation)
- Response variables were log-transformed: log(y + 1)

*Modeling approach*

We used confirmatory path analysis (Shipley, 2000, 2009). The piecewise structural equation models (SEM) were fitted using the R-package “piecewiseSEM” (Lefcheck, 2016). Indirect effects were calculated by multiplying standardized coefficients along pathways in the model. Confidence intervals for direct, indirect, and total effects for each predictor were calculated in the R-package “semEff” (Murphy, 2021) using nonparametric bootstrap with 1000 randomizations. Model coefficients are standardized coefficients.

*Response variables*

Taxonomic, phylogenetic, and functional diversity were estimated for multiple q-orders (0, 1, and 2). In addition to the observed hill diversity, we also estimated standardized coverage-based diversity (Chao et al., 2021).

- <u>tax_hill</u>: Taxonomic hill diversity
- <u>phy_hill</u>: Phylogenetic hill diversity
- <u>fun_hill</u>: Functional hill diversity

*Explanatory variables*

Predictors were arranged in four groups:

1. Experimental variables:
   - <u>planted_diversity</u>: The diversity of planted tree species (0, 1, 2, 3, 6)
   - <u>island_area</u>: Island area (m2) | Range: 25 m2 - 1600 m2

2. Soil properties:
   - <u>soil_pc1</u>: Associated with higher soil C, N, and lower bulk density
   - <u>soil_pc2</u>: Associated with higher soil P and pH

2. Vegetation structure:
   - <u>veg_pc1</u>: Associated with higher stand structural complexity
   - <u>veg_pc2</u>: Associated with higher tree dominance in the canopy and litter depth

2. Landscape context:
   - <u>forest_dist</u>: Distance to the nearest forest patch (m)
   - <u>iso_trees</u>: Number of isolated trees within a 100m radius

*Sem model specification and formula*

```
piecewiseSEM::psem(
    lm(hill ~ planted_diversity + island_area + soilPC1 + soilPC2 +
       vegPC1 + vegPC2 + Dist_forest + Iso_Trees, datset),
    lm(vegPC1  ~ planted_diversity, datset),
    lm(vegPC2  ~ island_area, datset)
  )
```

**Model validation**

- Model goodness of fit was tested with Fisher's C statistics (Shipley, 2000)
- Missing paths were evaluated with the d-separation test (Lefcheck, 2016)

For each linear model within SEM we also:

- Used a QQ-plot of residuals to check for normality
- Tested model residuals for spatial autocorrelation using Moran’s I coefficient
- Tested for homogeneity of variance
- Inspected residuals for multicollinearity
- Checked for influential observations (outliers) based on a composite outlier score (Lüdecke et al., 2021)

**Table S1.** Mechanistic framework and predictions linking the study variables with the diversity of regenerating woody species. Colors represent groups of drivers: experimental treatments, vegetation structure, soil properties, and landscape context.

| | Path | Relationship | Mechanism | Prediction |
|---|---|---|---|---|
| | **A** | Planted Tree diversity<br>⇩<br>Regenerating diversity | I- A more diverse internal source of seeds and vegetative reproduction<br><br>II - Different tree species modify environmental conditions and resource availability, promoting recruiting species with contrasting requirements for establishment and survival<br><br>II - Reduced competition due to complementarity between plant species and diluted pressure from natural enemies (i.e. herbivores, pathogens)<br><br>III- Diversity begets diversity: the higher richness of planted trees promotes small-scale heterogeneity in environmental conditions and complex, competitive networks that minimize species loss due to competitive exclusion | The diversity of the regenerating plant community increases with the richness of planted trees (Maynard et al., 2017; Palmer & Maurer, 1997; Paterno et al., 2016; Tinya et al., 2019). |
| | **B** | Island area<br>⇩<br>Regenerating diversity | I - Larger islands receive more seeds due to passive sampling (habitat quantity) and because dispersers are more likely to visit and spend more time in larger islands (habitat quality)<br><br>II - Larger islands are more heterogeneous and structurally complex (resources, abiotic & biotic environment)<br><br>III - Larger islands have reduced edge effects (buffer from oil palm plantations) | Diversity of the regenerating plant community increases with tree island area (Chase et al., 2020; Holl et al., 2020; MacArthur & Wilson, 1963; Zahawi & Augspurger, 2006; Zemp et al., 2023). |
| | **C** | Island area<br>⇩<br>Tree dominance | I - Trees grow faster in larger tree islands due to enhanced environmental heterogeneity, more resources, and reduced edge effects.<br><br>II - No thinning of oil-palm in smaller plots (direct effect of experimental design) | Larger island areas have higher tree cover and lower oil-palm dominance (Zemp, Gérard, et al., 2019; Zemp et al., 2023) |
| | **D** | Tree diversity<br>⇩<br>Vegetation complexity | I - Species with different functional traits and architecture promotes higher vegetation complexity<br><br>II - Higher niche complementarity promotes plant growth and thereby more occupation of 3D-space | Higher planted tree diversity increases vegetation complexity (Zemp, Ehbrecht, et al., 2019) |

| | Path | Relationship | Mechanism | Prediction |
|---|---|---|---|---|
| | **E** | Veg. structural Complexity<br>⇩<br>Regenerating diversity | I - Variable light, shade, temperature condition under more complex environments favors different regenerating niches<br>II - Higher complexity of the vegetation offer habitats for different seed dispersers (birds, bats, ants) | Higher structural complexity increases the diversity of regenerating woody species |
| | **F** | Tree dominance<br>⇩<br>Regenerating diversity | I - Higher tree dominance reduces understory cover of light-dependent species<br>II - Lower canopy cover of oil palms reduces competition and mortality of recruiting seedlings | Higher tree dominance increases the diversity of regenerating woody species. |
| | **G** | Isolated trees<br>⇩<br>Regenerating diversity | I - Scattered trees are an important source of seed propagules and stepping stones for seed dispersers | Tree islands surrounded by more isolated trees show higher diversity of naturally regenerating woody species (Guevara et al., 1986; Manning et al., 2006) |
| | **H** | Distance to forest<br>⇩<br>Regenerating diversity | I - Reduced isolation leads to higher colonization rate near forest patches<br>II - Higher amount of habitat in the surrounding landscape reduces dispersal limitation | Tree islands closer to forest patches show higher diversity of naturally regenerating trees (Fahrig, 2013; Crouzeilles & Curran, 2016) |
| | **I & J** | Soil Fertility & compaction<br>⇩<br>Regenerating diversity | I - Low nutrient availability (N, C, and P) limits seedling establishment<br>II - High soil compaction prevents seedling establishment due to physical barrier | Diversity of the regenerating plant community increases with increasing soil fertility and reduced soil compaction |

**Table S2.** List of recorded woody species – planted and regenerating – in EFForTS-BEE plots, Sumatra, Indonesia.

| Species | Family | Alien | Life form | Habitat | Dispersal |
|---|---|---|---|---|---|
| **Planted species** | | | | | |
| *Dyera polyphylla* (Miq.) Steenis | Apocynaceae | | tree | forest | |
| *Elaeis guineensis* Jacq. | Arecaceae | | tree | forest | |
| *Shorea leprosula* Miq. | Dipterocarpaceae | | tree | forest | |
| *Archidendron jiringa* (Jack) I.C.Nielsen | Fabaceae | | tree | forest | |
| *Parkia speciosa* Hassk. | Fabaceae | | tree | forest | |
| *Peronema canescens* Jack | Lamiaceae | | tree | forest | |
| *Durio zibethinus* L. | Malvaceae | | tree | forest | |
| **Regenerating species** | | | | | |
| *Alstonia angustiloba* Miq. | Apocynaceae | | tree | forest | anemochory |
| *Tabernaemontana pauciflora* Blume | Apocynaceae | | treelet | forest | zoochory |
| *Willughbeia* sp.DMF0064 | Apocynaceae | | liana | | zoochory |
| *Elaeis guineensis* Jacq. | Arecaceae | yes | tree | | zoochory |
| *Clibadium surinamense* L. | Asteraceae | yes | shrub | open | zoochory |
| *Trema tomentosum* (Roxb.) H.Hara | Cannabaceae | | tree | open | zoochory |
| *Terminalia* cf. *calamansanai* (Blanco) Rolfe | Combretaceae | | tree | open | zoochory |
| *Camonea pilosa* (Houtt.) A.R.Simões & Staples | Convolvulaceae | | liana | open | zoochory |
| *Erycibe rheedei* Blume | Convolvulaceae | | liana | open | zoochory |
| *Dillenia* cf. *excelsa* (Jack) Martelli ex Gilg. | Dilleniaceae | | tree | forest | zoochory |
| *Croton argyratus* Blume | Euphorbiaceae | | tree | forest | anemochory |
| *Macaranga bancana* (Miq.) Müll.Arg. | Euphorbiaceae | | tree | open | zoochory |
| *Macaranga conifera* (Rchb.f. & Zoll.) Müll.Arg. | Euphorbiaceae | | tree | forest | zoochory |
| *Macaranga gigantea* (Rchb.f. & Zoll.) Müll.Arg. | Euphorbiaceae | | tree | open | zoochory |
| *Macaranga hosei* King ex Hook.f. | Euphorbiaceae | | tree | open | zoochory |
| *Macaranga trichocarpa* (Zoll.) Müll.Arg. | Euphorbiaceae | | shrub | open | zoochory |
| *Mallotus macrostachyus* (Miq.) Müll.Arg. | Euphorbiaceae | | tree | open | zoochory |
| *Mallotus paniculatus* (Lam.) Müll.Arg. var. *paniculatu*s | Euphorbiaceae | | tree | open | zoochory |
| *Mallotus peltatus* (Geiseler) Müll.Arg. | Euphorbiaceae | | tree | open | zoochory |
| *Archidendron jiringa* (Jack) I.C.Nielsen | Fabaceae | | tree | forest | zoochory |

| Species | Family | Alien | Life form | Habitat | Dispersal |
|---|---|---|---|---|---|
| *Centrosema molle* Mart. ex Benth. | Fabaceae | yes | liana | open | zoochory |
| *Mucuna biplicata* Teijsm. & Binn. ex Kurz | Fabaceae | | liana | open | anemochory |
| *Phanera semibifida* (Roxb.) Benh. var. *semibifida* | Fabaceae | | liana | open | zoochory |
| *Senna alata* (L.) Roxb. | Fabaceae | yes | shrub | open | anemochory |
| *Cratoxylum formosum* (Jack) Benth. & Hook.f. ex Dyer | Hypericaceae | | tree | forest | anemochory |
| *Callicarpa pentandra* Roxb. | Lamiaceae | | tree | open | zoochory |
| *Clerodendrum disparifolium* Blume | Lamiaceae | | treelet | forest | zoochory |
| *Peronema canescen*s Jack | Lamiaceae | | tree | open | anemochory |
| *Vitex quinata* (Lour.) F.N.Williams | Lamiaceae | | tree | forest | zoochory |
| *Litsea umbellata* (Lour.) Merr. | Lauraceae | | tree | open | zoochory |
| *Commersonia bartramia* (L.) Merr. | Malvaceae | | tree | open | zoochory |
| *Hibiscus macrophyllus* Roxb. ex Hornem. | Malvaceae | | tree | open | anemochory |
| *Trichospermum javanicum* Blume | Malvaceae | | tree | forest | anemochory |
| *Urena lobata* L. ssp. *sinuat*a | Malvaceae | | shrub | open | zoochory |
| *Bellucia pentamer*a Naudin | Melastomataceae | yes | tree | open | zoochory |
| *Melastoma malabathricum* L. ssp. *malabathricum* | Melastomataceae | | shrub | open | zoochory |
| *Swietenia macrophylla* King | Meliaceae | yes | tree | forest | anemochory |
| *Ficus aurata* (Miq.) Miq. | Moraceae | | treelet | open | zoochory |
| *Ficus caulocarpa* (Miq.) Miq. | Moraceae | | tree | forest | zoochory |
| *Ficus glandulifera* (Wall. ex Miq.) King | Moraceae | | tree | forest | zoochory |
| *Ficus padana* Burm.f. | Moraceae | | tree | open | zoochory |
| *Ficus variegata* Blume | Moraceae | | tree | open | zoochory |
| *Ficus vrieseana* Miq. | Moraceae | | tree | open | zoochory |
| *Sloetia elongata* (Miq.) Koord. | Moraceae | | tree | open | zoochory |
| *Champereia manillana* (Blume) Merr. | Opiliaceae | | treelet | forest | zoochory |
| *Galearia filiformis* (Blume) Boerl. | Pandaceae | | tree | forest | zoochory |
| *Eurya nitida* Korth. | Pentaphylacaceae | | treelet | open | zoochory |
| *Breynia racemosa* (Blume) Müll.Arg. | Phyllanthaceae | | treelet | open | zoochory |
| *Bridelia glauca* Blume var. *glauca* | Phyllanthaceae | | tree | forest | zoochory |
| *Glochidion borneense* (Müll.Arg.) Boerl. | Phyllanthaceae | | tree | forest | zoochory |
| *Gigantochloa scortechinii* Gamble | Poaceae | | shrub | open | zoochory |

| Species | Family | Alien | Life form | Habitat | Dispersal |
|---|---|---|---|---|---|
| *Canthium horridum* Blume | Rubiaceae | | shrub | open | zoochory |
| *Mussaenda frondosa* L. | Rubiaceae | | liana | open | zoochory |
| *Neolamarckia cadamba* (Roxb.) Bosser | Rubiaceae | | tree | | zoochory |
| *Neonauclea calycina* (Bartl. ex DC.) Merr. | Rubiaceae | | tree | open | zoochory |
| *Uncaria cordata* (Lour.) Merr. | Rubiaceae | | liana | open | anemochory |
| *Urophyllum peltistigma* Miq. | Rubiaceae | | tree | forest | zoochory |
| *Clausena excavata* Burm.f. | Rutaceae | | tree | open | zoochory |
| *Homalium caryophyllaceum* (Zoll. & Moritzi) Benth. | Salicaceae | | treelet | forest | zoochory |
| *Nephelium* sp. | Sapindaceae | | tree | | zoochory |
| *Brucea javanica* (L.) Merr. | Simaroubaceae | | treelet | open | zoochory |
| *Solanum jamaicense* Mill. | Solanaceae | yes | shrub | open | zoochory |
| *Stachytarpheta australis* Moldenke | Verbenaceae | yes | shrub | open | zoochory |
| *Leea indica* (Burm.f.) Merr. | Vitaceae | | treelet | open | zoochory |
| climber DMF0061 | unknown | | liana | | zoochory |

**Table S3.** Results of linear models between observed alpha plant diversity (q=1, taxonomic, phylogenetic, and functional) against tree island area (log) and restoration treatment.

| | **Taxonomic** | | **Phylogenetic** | | **Functional** | |
|---|---|---|---|---|---|---|
| *Predictors* | *Estimates* | *CI* | *Estimates* | *CI* | *Estimates* | *CI* |
| (Intercept) | -1.01 *** | -1.46 – -0.56 | -0.79 *** | -1.14 – -0.43 | -0.46 ** | -0.73 – -0.19 |
| log (island area) | 0.22 *** | 0.16 – 0.28 | 0.17 *** | 0.12 – 0.22 | 0.10 *** | 0.06 – 0.14 |
| Oil palm | *Reference* | | *Reference* | | *Reference* | |
| No planting | 1.12 *** | 0.63 – 1.62 | 1.06 *** | 0.66 – 1.45 | 0.89 *** | 0.59 – 1.19 |
| Mono | 1.28 *** | 0.90 – 1.66 | 1.18 *** | 0.88 – 1.48 | 0.95 *** | 0.73 – 1.18 |
| Mixture | 1.50 *** | 1.12 – 1.87 | 1.31 *** | 1.01 – 1.61 | 1.12 *** | 0.90 – 1.35 |
| Observations | 56 | | 56 | | 56 | |
| $R^2$ / $R^2$ adjusted | 0.72 / 0.70 | | 0.74 / 0.72 | | 0.74 / 0.72 | |

* *p<0.05* ** *p<0.01* *** *p<0.001*

**Table S4.** Results from linear models predicting observed and standardized taxonomic, phylogenetic, and functional diversity of the regenerating plant community for q-order 1.

| | **Observed** | | | **Standardized** | | |
|---|---|---|---|---|---|---|
| | **Taxonomic** | **Phylogenetic** | **Functional** | **Taxonomic** | **Phylogenetic** | **Functional** |
| *Predictors* | *Estimates* | *Estimates* | *Estimates* | *Estimates* | *Estimates* | *Estimates* |
| (Intercept) | **1.52** *** | **1.35** *** | **1.10** *** | **1.28** *** | **1.17** *** | **1.00** *** |
| Veg. PC1 | -0.02 | -0.03 | 0.00 | 0.02 | -0.00 | 0.01 |
| Veg. PC2 | **0.14** * | **0.11** * | 0.05 | **0.22** ** | **0.17** ** | **0.09** * |
| Soil PC1 | **0.15** ** | **0.11** ** | **0.08** ** | **0.14** ** | **0.11** * | **0.08** * |
| Soil PC2 | 0.05 | 0.04 | 0.04 | 0.09 | 0.08 | 0.06 |
| Tree Diversity | **0.11** * | 0.07 | **0.07** * | 0.09 | 0.07 | **0.07** * |
| Island area | **0.20** ** | **0.16** ** | **0.09** * | 0.05 | 0.05 | 0.03 |
| Dist. Forest | 0.01 | 0.02 | 0.04 | -0.00 | 0.00 | 0.02 |
| Isolated trees | 0.03 | 0.02 | 0.02 | -0.01 | -0.01 | -0.00 |
| Observations | 52 | 52 | 52 | 52 | 52 | 52 |
| $R^2$ | 0.68 | 0.65 | 0.60 | 0.55 | 0.54 | 0.51 |

* *p<0.05* ** *p<0.01* *** *p<0.001*

**Table S5.** List of predictors used in the piecewise structural equation models.

| Group | Variable | Description | Unit |
|---|---|---|---|
| Experimental | Planted tree diversity | The number of tree species planted in each plot (0, 1, 2, 3, and 6) | |
| Experimental | Island area | The total area of the plot (25 $m^2$, 100 $m^2$, 400 $m^2$, 1600 $m^2$) | $m^2$ |
| Soil properties | Soil PC1 | The first axis of the PCA analysis with 7 soil variables. See Figure S2. | |
| Soil properties | Soil PC2 | The second axis of the PCA analysis with 7 soil variables. See Figure S2. | |
| Vegetation structure | Veg PC1 | The first component of the PCA analysis represents a gradient from open to dense and complex vegetation ("structural complexity"). See Figure S2. | |
| Vegetation structure | Veg -PC2 | The second component of the PCA analysis represents a gradient from oil palm to tree dominance ("tree dominance). See Figure S2. | |
| Landscape context | Distance to the forest | The distance to the nearest forest patch within a 100 m radius. The minimum patch size was 0.5 hectares. | m |
| Landscape context | Number of isolated trees | The number of trees outside of forest patches (isolated) within a 100 radius of tree islands | |

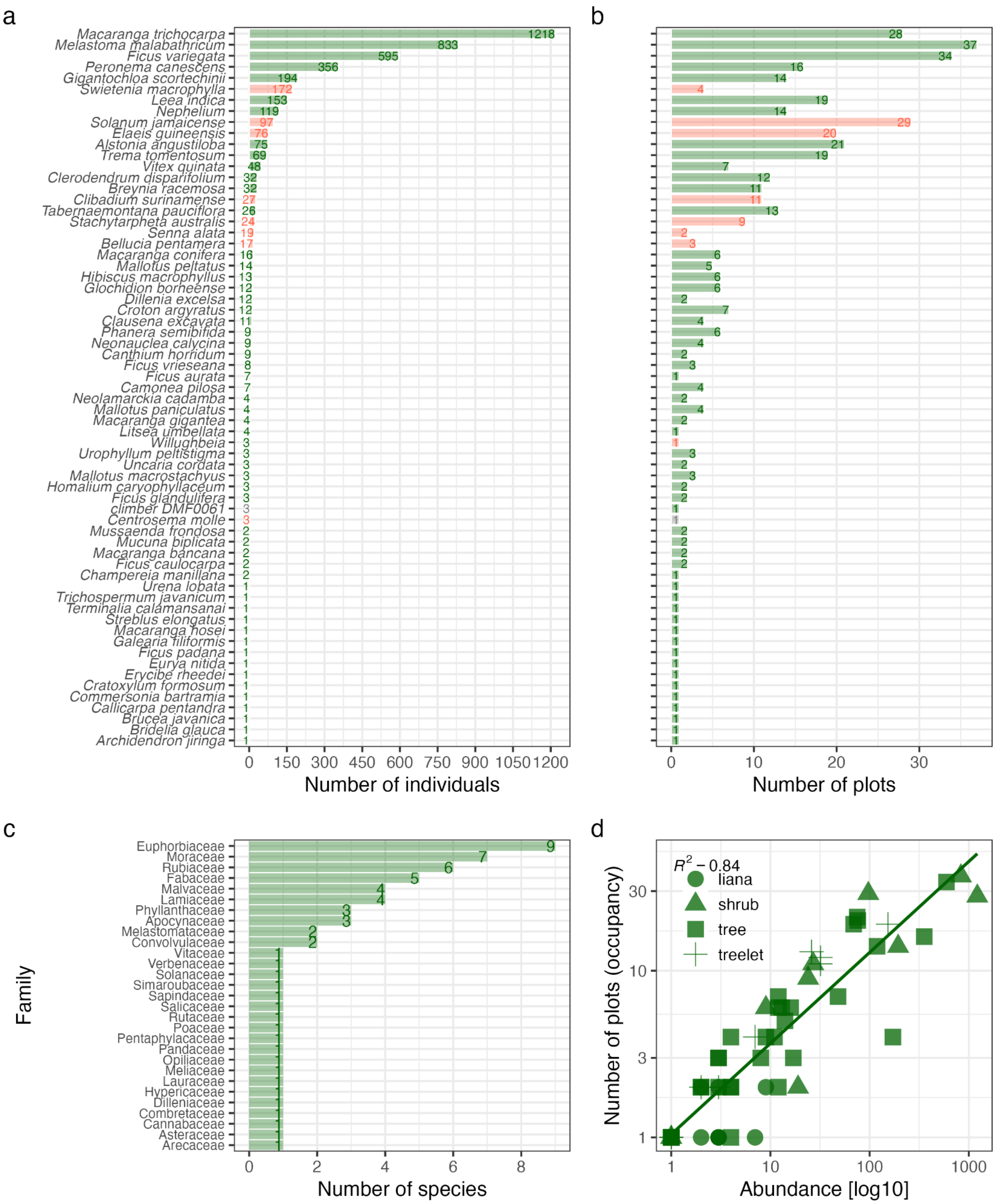

**Figure S1.** Taxonomic overview of regenerating woody species in EFForTS-BEE experiment. Number of individuals per species (a), number of plots where species registered (occupancy) (b), number of species per family (c), and the relationship between total abundance and species occupancy (proportion of plots) (d). Red bars represent species alien to Sumatra, Indonesia.

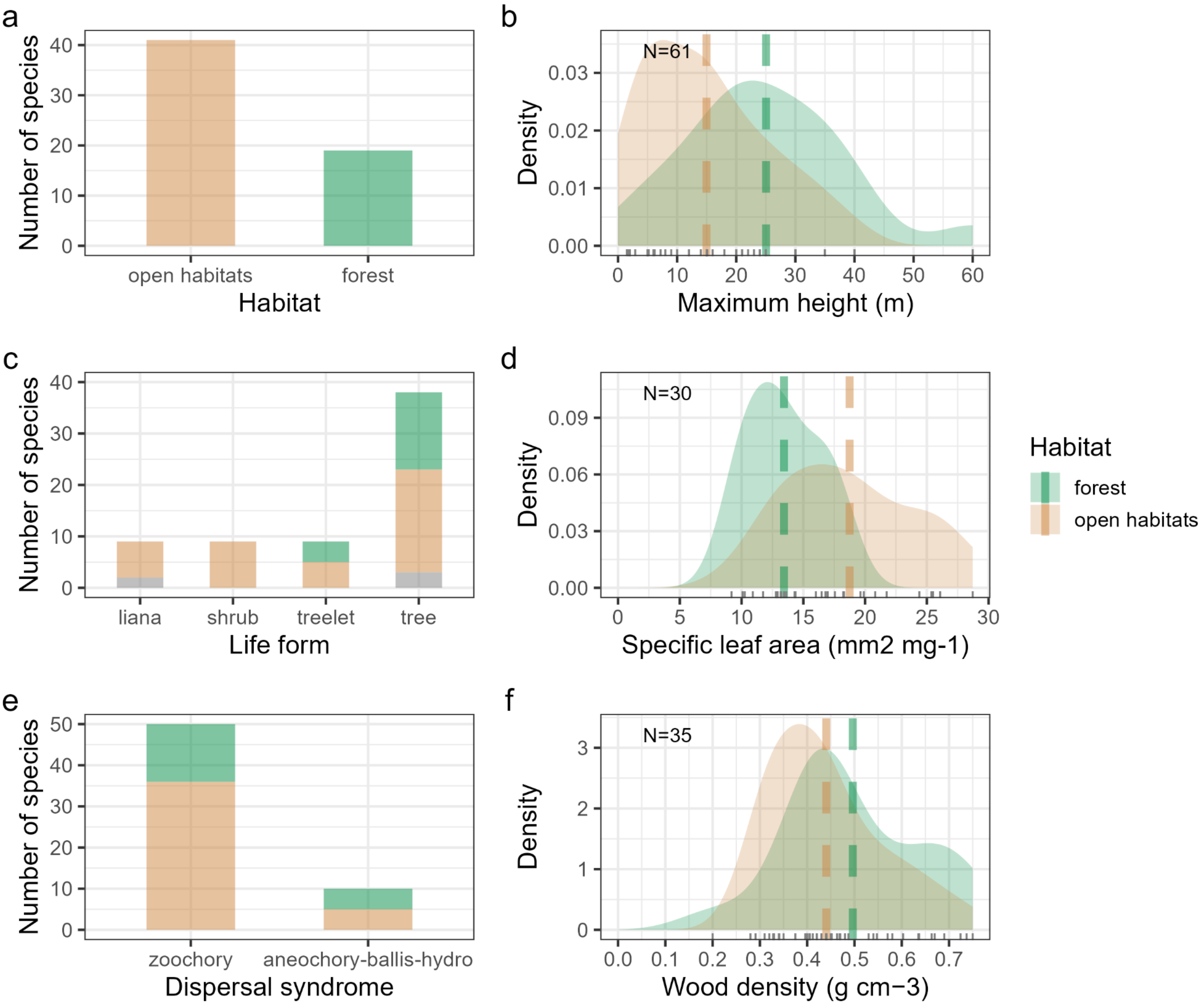

**Figure S2.** Ecological strategy across regenerating woody species in the EFForTS-BEE experiment, Sumatra, Indonesia. Distribution of habitat (a), specific leaf area (b), life form (c), specific leaf area (d), dispersal syndrome (e), and wood density (f). The number of species with data is shown in the top left corner of each plot. The dashed green line represents the median value across species. In plot (e), zoochory includes endozoochory, epizoochory, and myrmecochory. Plots are collared by habitat type. Gray colors represent missing data.

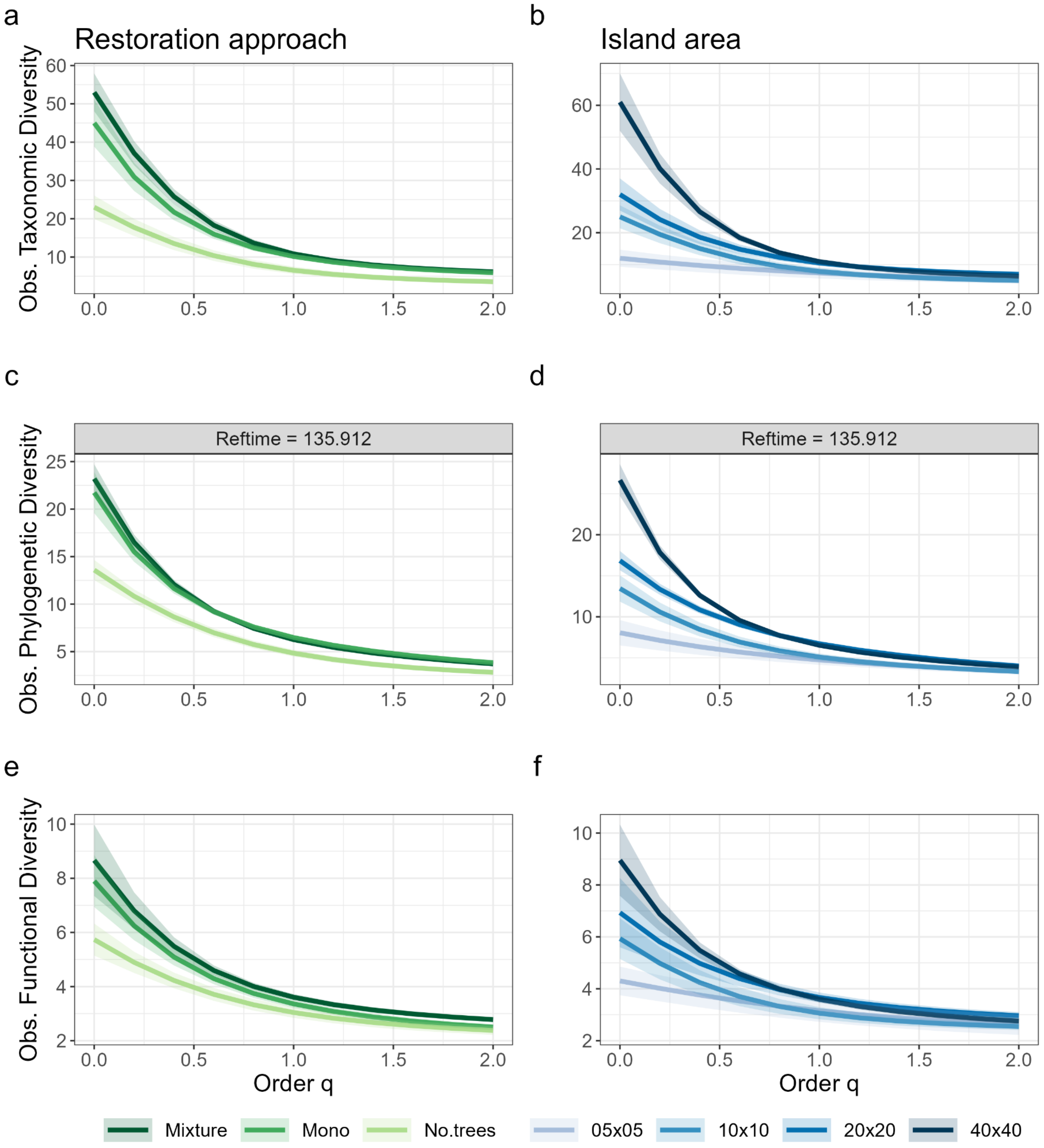

**Figure S3. Diversity profiles across restoration approach and island area.** Observed taxonomic (a,b), phylogenetic (c, d), and functional (e, f) Hill diversity across different restoration treatments and tree island sizes. Hill diversity is plotted against a continuous gradient of q-orders, ranging from q=0 (species richness), to q=2 (inverse of the Simpson index). Solid lines represent average values and shaded areas represent 95% bootstrap confidence intervals (N = 100).

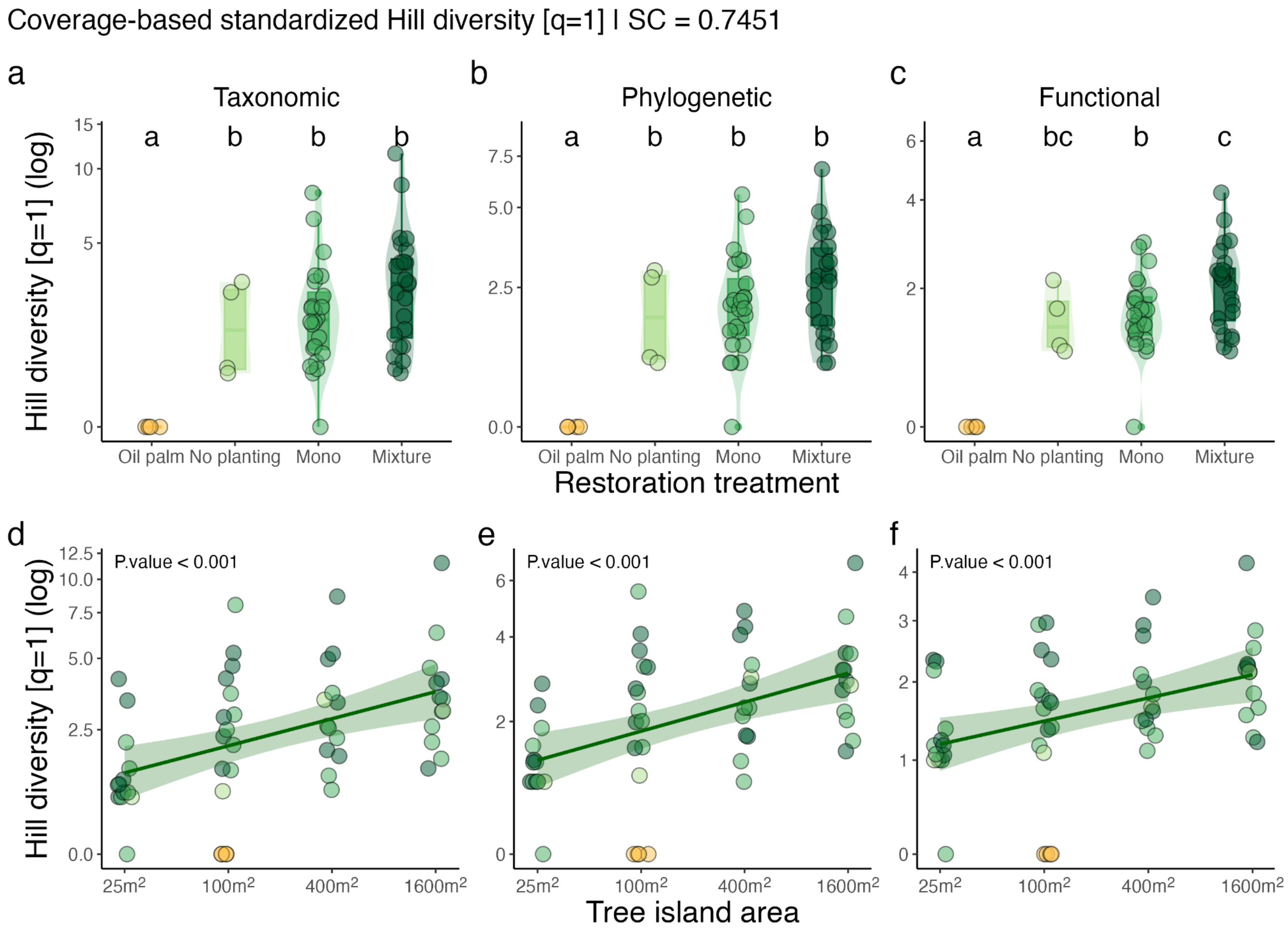

**Figure S4. Experimental effects on standardized multifaceted alpha diversity of recruiting species**. Relationship between coverage-based standardized taxonomic, phylogenetic, and functional diversity (q=1) against restoration treatment (a, b, c) and plot area (d, e, f). Yellow points represent Conventional Oil Pal monoculture, while green points represent tree islands. Darker green represents higher planted tree diversity, and light green represents plots with no planted trees. Letters represent pairwise comparisons across treatment levels through Least-squares means with p-values adjusted by the Tukey method. Diversity estimates were standardized to the minimum sample coverage (SC) among treatments extrapolated to double reference size (Chao et al. 2021).

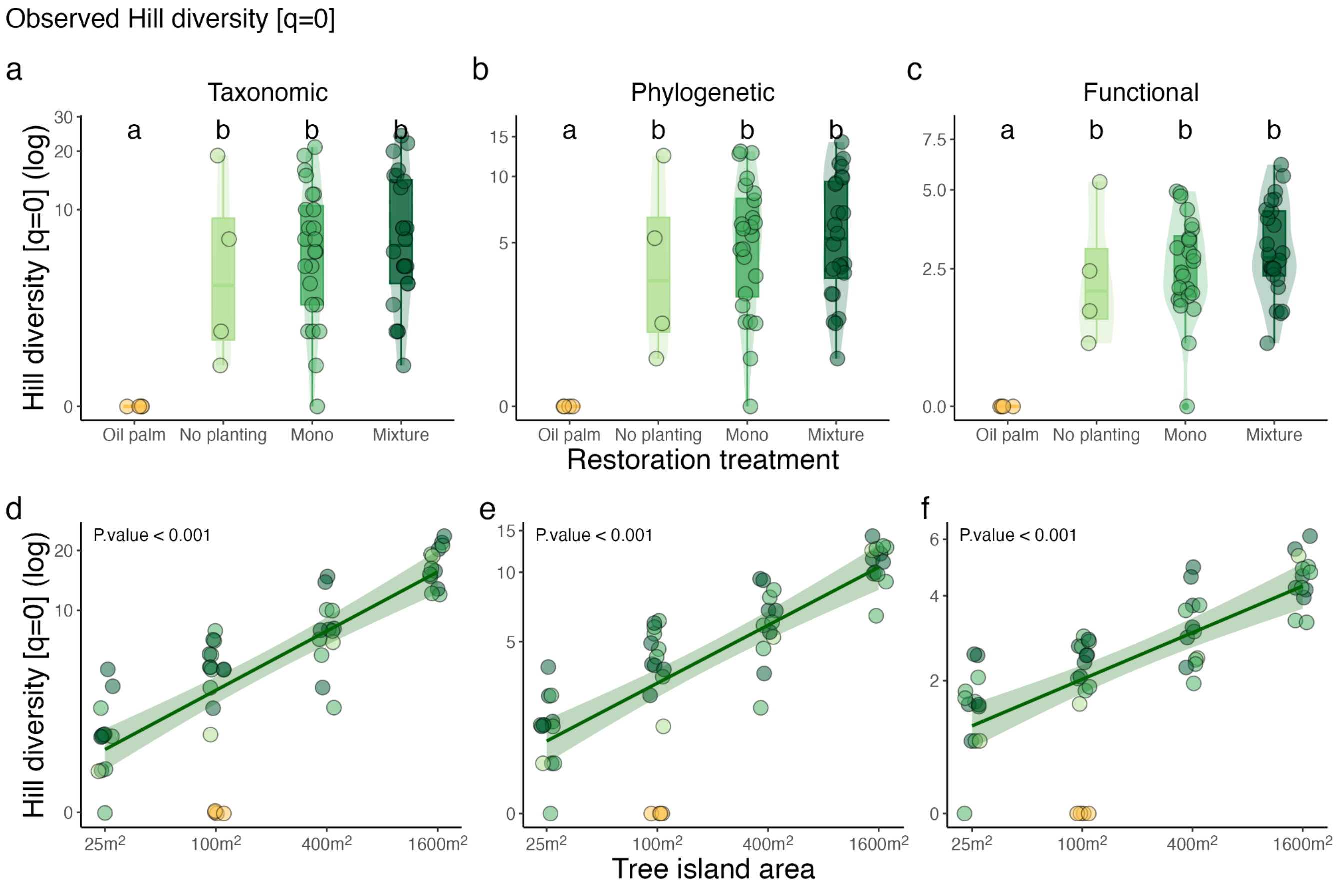

**Figure S5. Experimental effects on observed multifaceted alpha diversity [q=0] of recruiting species**. Relationship between coverage-based standardized taxonomic, phylogenetic, and functional diversity (q=0) against restoration treatment (a, b, c) and plot area (d, e, f). Yellow points represent Conventional Oil Pal monoculture, while green points represent tree islands. Darker green represents higher planted tree diversity, and light green represents plots with no planted trees. Letters represent pairwise comparisons across treatment levels through Least-squares means with p-values adjusted by the Tukey method.

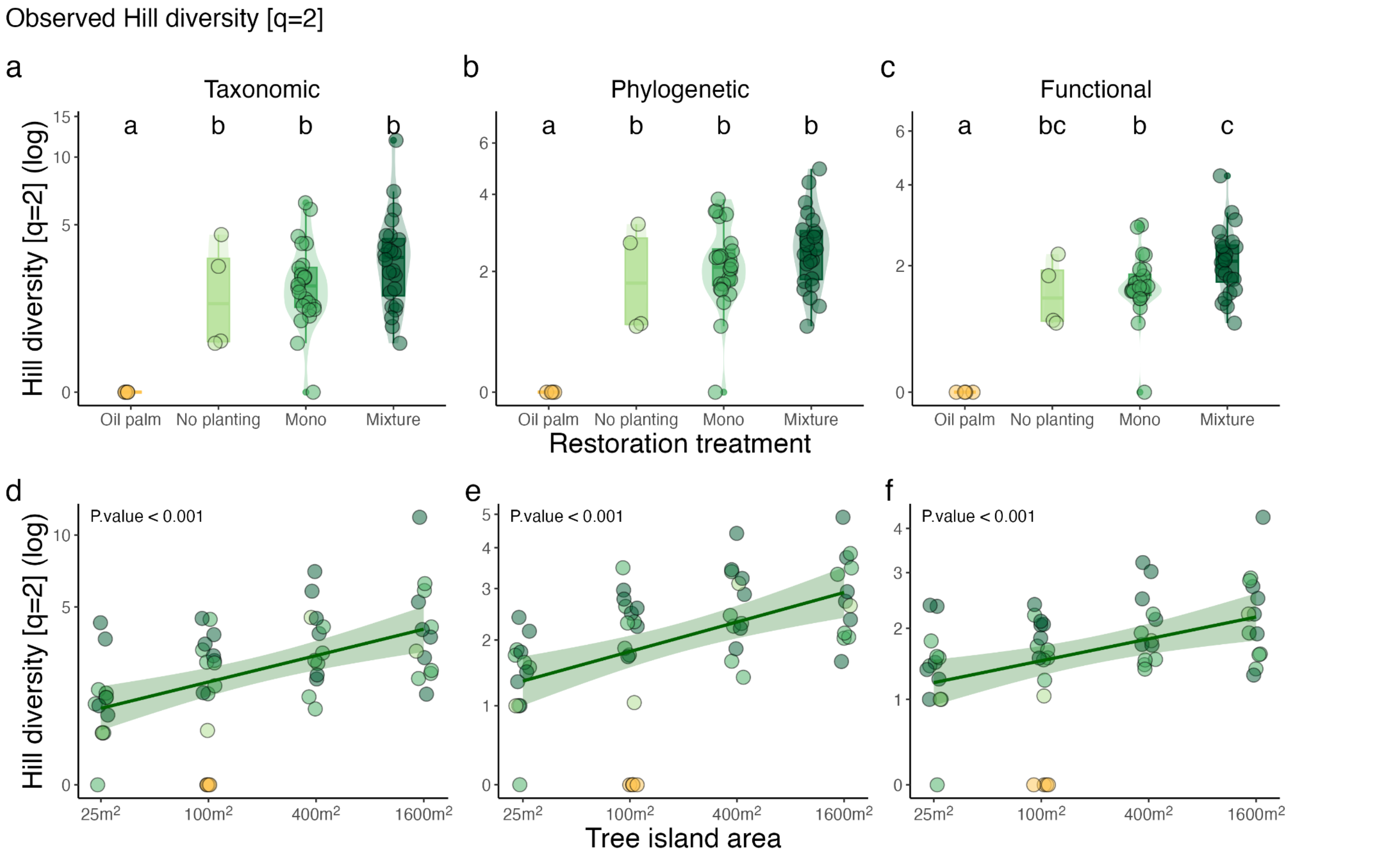

**Figure S6. Experimental effects on observed multifaceted alpha diversity [q=2] of recruiting species**. Relationship between coverage-based standardized taxonomic, phylogenetic, and functional diversity (q=2) against restoration treatment (a, b, c) and plot area (d, e, f). Yellow points represent Conventional Oil Pal monoculture, while green points represent tree islands. Darker green represents higher planted tree diversity, and light green represents plots with no planted trees. Letters represent pairwise comparisons across treatment levels through Least-squares means with p-values adjusted by the Tukey method.

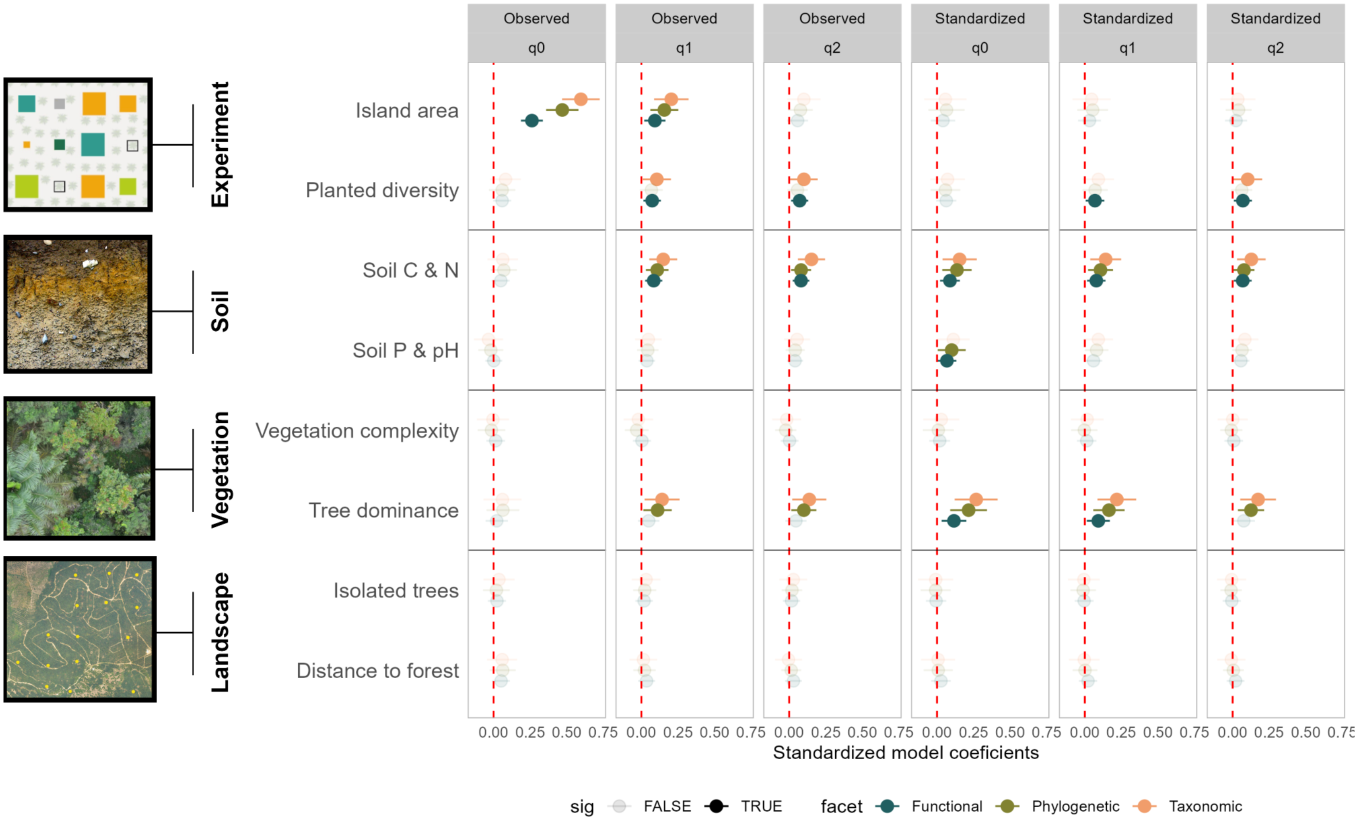

**Figure S7. Results from Piecewise Structural Equation Models.** Standardized model coefficients from SEM for observed (first three columns) and standardized (last three columns) hill diversity of q-order 0, 1, and 2. Shaded dots represent non-significant predictors ($p > 0.05$). Predictors were grouped into experimental treatment (island area and planted tree diversity), soil properties (soil PC1 and soil PC2), vegetation structure (vegetation PC1 and -PC2), and landscape context (number of isolated trees and distance to the nearest forest).

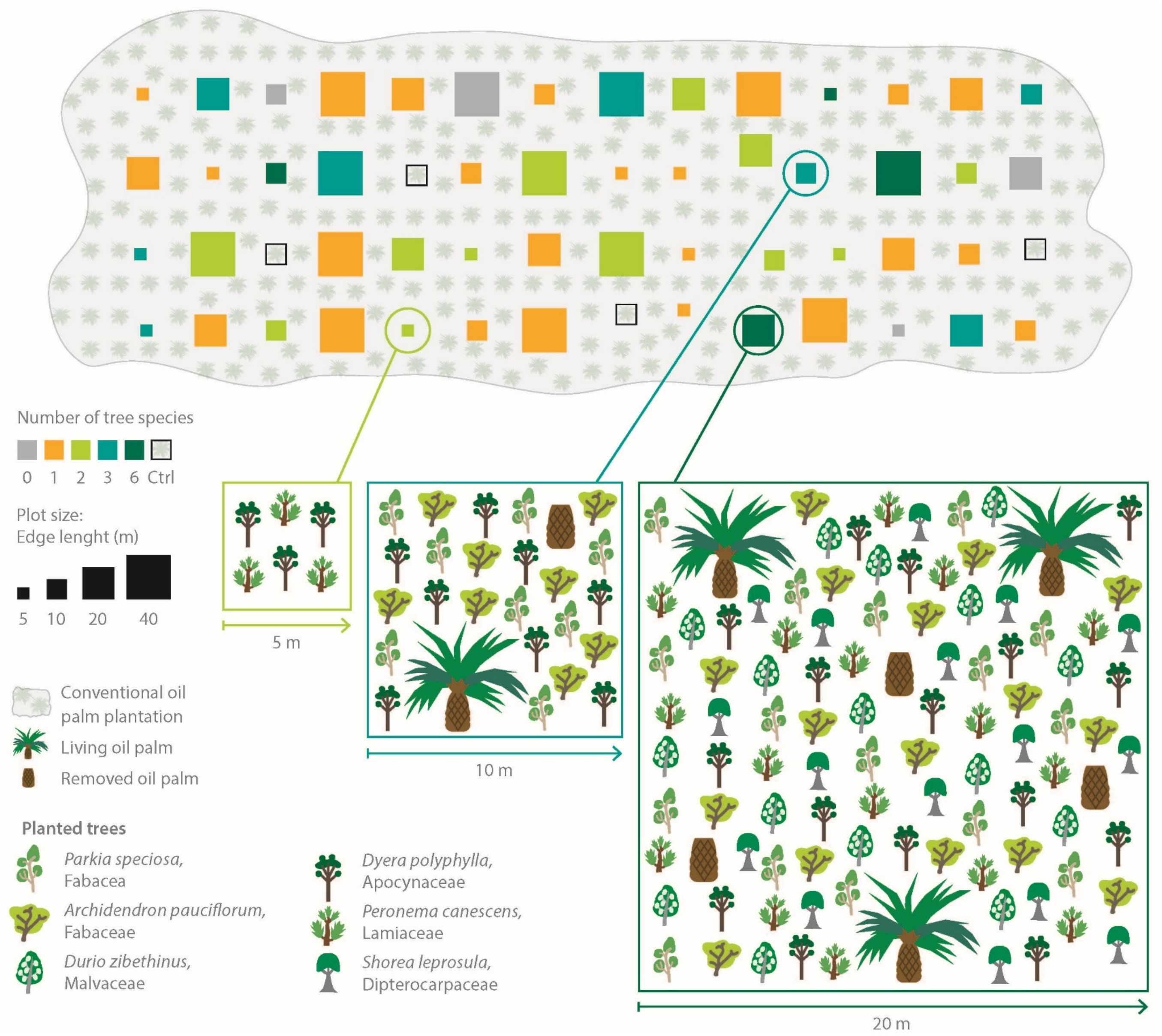

**Figure S8. EFForTS-BEE experimental design**. The top panel shows the distribution of plots within an industrial oil palm plantation. Different colors represent diversity levels (0, 1, 2, 3, 6), and gray squares are control plots (conventional oil palm plantation). Squares with different sizes represent the experimental treatment of varying plot sizes (5x5m, 10x10m, 20x20m, 40x40m). The lower panel shows the spatial arrangement of planted trees, oil palms, and oil palms that were removed. See Teuscher et al. (2016) for the details.

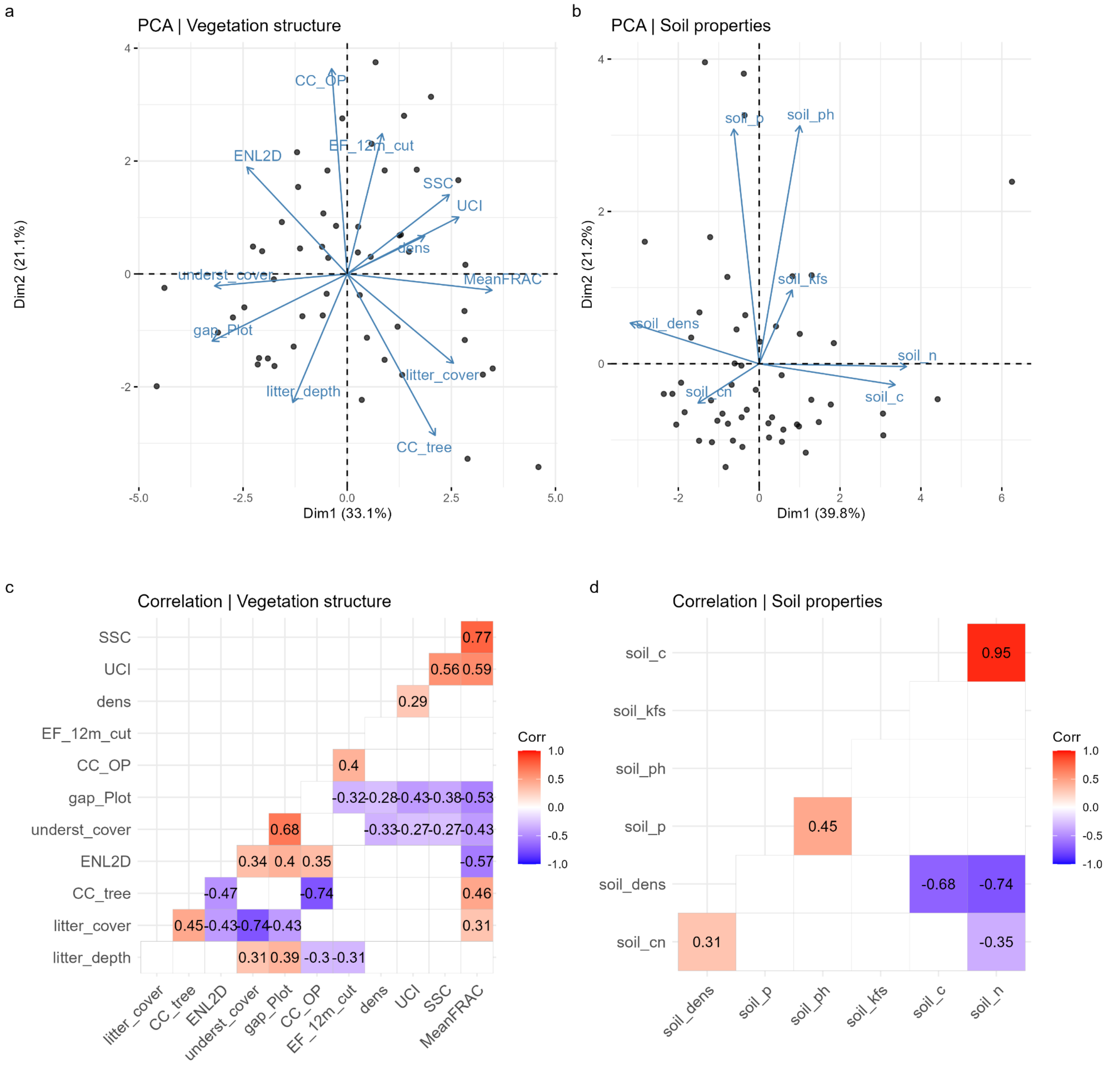

**Figure S9. Principal component Analysis and correlation matrix for vegetation structure and soil properties**. Definition of variables for the PCA on vegetation structure (a): (1) stand structural complexity index (SSC), (2) the mean fractal dimension index (MeanFRAC), (3) the effective number of layers (ENL) and (4) the understory complexity index (UCI). Using hemispherical photo and drone-based photogrammetry (Khotkhong et al. 2019), we calculated (5) canopy gap fraction that was partitioned as (6) oil palm cover and (7) tree cover; and (8) oil palm density. We conducted ground-based assessment of (9) tree density, (10) understorey vegetation cover, (11) litter cover, and (12) litter depth. More information on the structural variables and the PCA can be found in Zemp et al. 2023. Definition of variables for the PCA on soil properties (b).

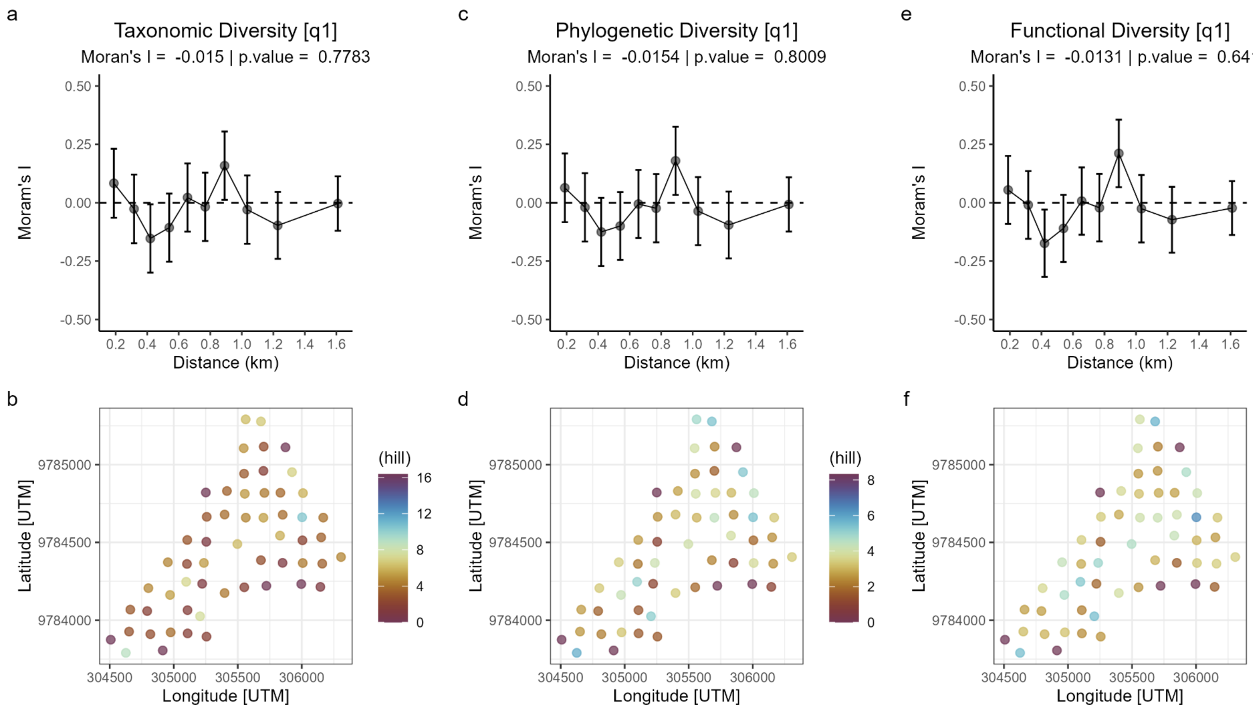

**Figure S10.** Spatial correlograms based on Moran's I index for taxonomic (a), phylogenetic (c), and functional hill diversity of q-order = 0 (e). The lower row of plots shows the spatial distribution of hill diversity across all 56 experimental plots. Hill diversity was log-transformed before analysis [log(x + 1)]. Global Moran’s I test is provided on the top of the plots.

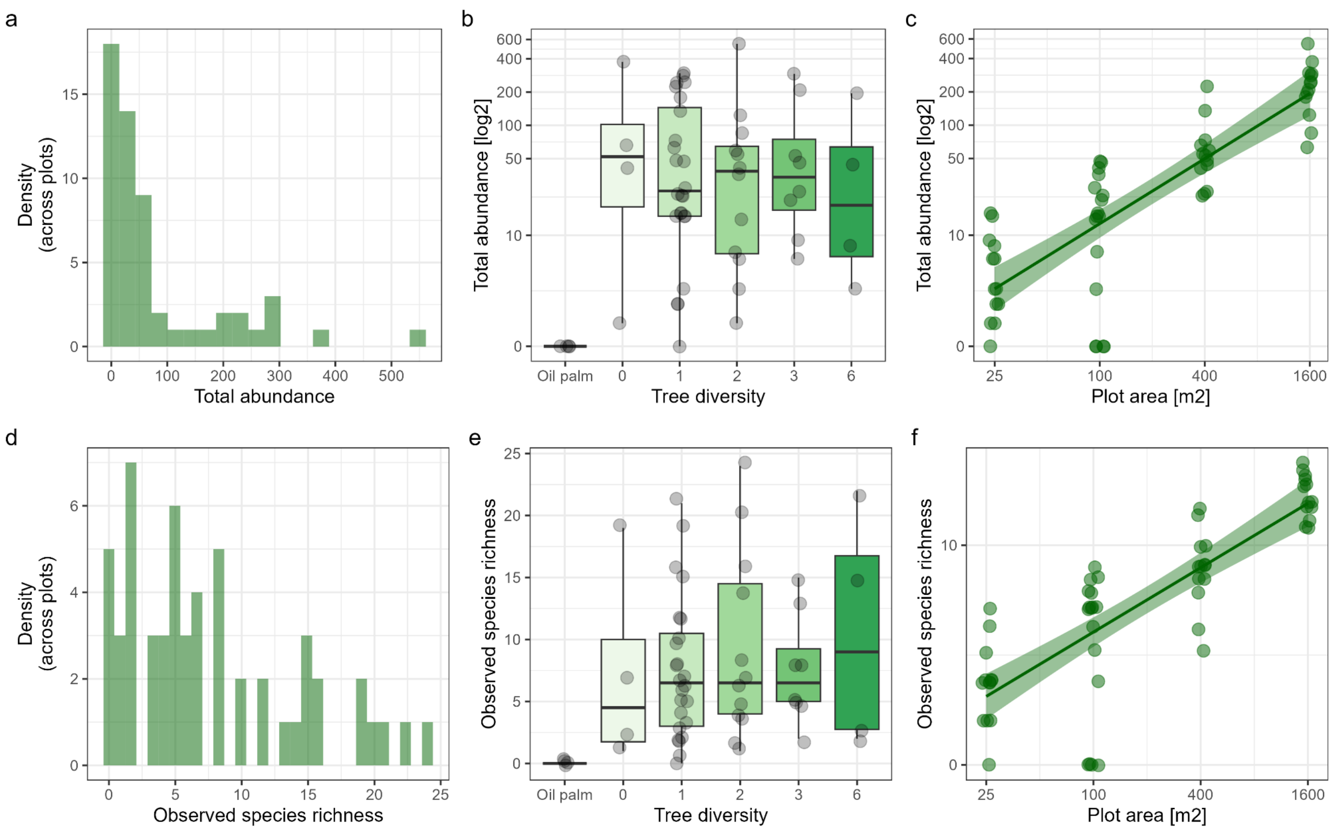

**Figure S11. Abundance and observed richness of woody plant species regenerating in the EFForTS-BEE experiment.** Distribution of total abundance (a) and observed species richness (d) across plots. Total abundance (b, c) and observed species richness (e, f) across tree diversity levels and plot area. Abundance and plot area were log-transformed (base 2) before analysis.
